# Optical control of ERK and AKT signaling promotes axon regeneration and functional recovery of PNS and CNS in *Drosophila*

**DOI:** 10.1101/2020.05.15.098251

**Authors:** Qin Wang, Huaxun Fan, Feng Li, Savanna S. Skeeters, Vishnu Krishnamurthy, Yuanquan Song, Kai Zhang

## Abstract

Neuroregeneration is a dynamic process synergizing the functional outcomes of multiple signaling circuits. Channelrhodopsin-based optogenetics shows feasibility of stimulating neural repair but does not pin down specific signaling cascades. Here, we utilized optogenetic systems, optoRaf and optoAKT, to delineate the contribution of the ERK and AKT signaling pathways to neuroregeneration in live *Drosophila* larvae. We showed that optoRaf or optoAKT activation not only enhanced axon regeneration in both regeneration competent and incompetent sensory neurons in the peripheral nervous system, but also allowed temporal tuning and proper guidance of axon regrowth. Furthermore, optoRaf and optoAKT differ in their signaling kinetics during regeneration, showing a gated versus graded response, respectively. Importantly in the central nervous system, their activation promotes axon regrowth and functional recovery of the thermonociceptive behavior. We conclude that non-neuronal optogenetics target damaged neurons and signaling subcircuits, providing a novel strategy in the intervention of neural damage with improved precision.

## Introduction

Inadequate neuroregeneration remains a major roadblock towards functional recovery after nervous system damage such as stroke, spinal cord injury (SCI), and multiple sclerosis. Extracellular factors from oligodendrocyte, astroglial, and fibroblastic sources restrict axon regrowth (Liu et al. 2006; Yiu and He 2006; Liu et al. 2011; Lu et al. 2014; Schwab and Strittmatter 2014) but eliminating these molecules only allows limited sprouting (Sun and He 2010), suggesting a down-regulation of the intrinsic regenerative program in injured neurons (Sun and He 2010; He and Jin 2016). The neurotrophic signaling pathway, which regulates neurogenesis during embryonic development, represents an important intrinsic regenerative machinery (Ramer et al. 2000). For instance, elimination of the PTEN phosphatase, an endogenous brake for neurotrophic signaling, yields axonal regeneration (Park et al. 2008).

An important feature of the neurotrophin signaling pathway is that the functional outcome depends on signaling kinetics (Marshall 1995) and subcellular localization (Watson et al. 2001). Indeed, neural regeneration from damaged neurons is synergistically regulated by multiple signaling circuits in space and time. However, pharmacological and genetic approaches do not provide sufficient spatial and temporal resolutions in the modulation of signaling outcomes in terminally differentiated neurons *in vivo*. Thus, the functional link between signaling kinetics and functional recovery of damaged neurons remains unclear. The emerging non-neuronal optogenetic technology uses light to control protein-protein interaction and enables light-mediated signaling modulation in live cells and multicellular organisms (Zhang and Cui 2015; Khamo et al. 2017; Johnson and Toettcher 2018; Leopold et al. 2018; Dagliyan and Hahn 2019; Goglia and Toettcher 2019). By engineering signaling components with photoactivatable proteins, one can use light to control a number of cellular processes, such as gene transcription (Motta-Mena et al. 2014; Wang et al. 2017), phase transition (Shin et al. 2017; Dine et al. 2018), cell motility (Wu et al. 2009) and differentiation (Khamo et al. 2019), ion flow across membranes (Kyung et al. 2015; Ma et al. 2018), and metabolism (Zhao et al. 2018; Zhao et al. 2019), to name a few. We have previously developed optogenetic systems named optoRaf (Zhang et al. 2014; Krishnamurthy et al. 2016) and optoAKT (Ong et al. 2016), which allow for precise control of the Raf/MEK/ERK and AKT signaling pathways, respectively. We demonstrated that timed activation of optoRaf enables functional delineation of ERK activity in mesodermal cell fate determination during *Xenopus laevis* embryonic development (Krishnamurthy et al. 2016). However, it remains unclear if spatially localized, optogenetic activation of ERK and AKT activity allows for subcellular control of cellular outcomes.

In this study, we used optoRaf and optoAKT to specifically activate the Raf/MEK/ERK and AKT signaling subcircuits, respectively. We found that both optoRaf and optoAKT activity enhanced axon regeneration in the regeneration-potent class IV da (C4da) and the regeneration-incompetent class III da (C3da) sensory neurons in *Drosophila* larvae, although optoRaf but not optoAKT enhanced dendritic branching. Temporally programmed and spatially restricted light stimulation showed that optoRaf and optoAKT differ in their signaling kinetics during regeneration and that both allow spatially guided axon regrowth. Furthermore, using a thermonociception based behavioral recovery assay, we found that optoRaf and optoAKT activation led to effective axon regeneration as well as functional recovery after central nervous system (CNS) injury. We note that most of previous optogenetic control of neural repair studies were based on channelrhodopsion in *C. elegens* (Sun et al. 2014), mouse DRG culture (Park et al. 2015a) or motor neuron-schwann cell co-culture (Hyung et al. 2019). Another study used blue-light activatable adenylyl cyclase bPAC to stimulate neural repair in mouse refractory axons (Xiao et al. 2015). These work highlights the feasibility of using optogenetics to study neural repair but did not pin down the exact downstream signaling cascade mediating neuronal repair. Additionally, most studies focused on peripheral neurons that are endogenously regenerative. Here, we specifically activated the ERK and AKT signaling pathways and performed a comprehensive study of neural regeratnion in both peripheral nervous system (PNS) and CNS neurons in live *Drosophila*. We envision that features provided by non-neuronal optogenetics, including reversibility, functional delineation, and spatiotemporal control will lead to a better understanding of the link between signaling kinetics and functional outcome of neurotrophic signaling pathways during neuroregeneration.

## Results

### Light enables reversible activation of the Raf/MEK/ERK and AKT signaling pathways

To reversibly control the Raf/MEK/ERK and AKT signaling pathways, we constructed a single-transcript optogenetic system using the p2A bicistronic construct that co-expresses fusion-proteins with the N-terminus of cryptochrome-interacting basic-helix-loop-helix (CIBN) and the photolyase homology region of cryptochrome 2 (CRY2PHR, abbreviated as CRY2 in this work). Following a similar design of the optimized optoRaf (Krishnamurthy et al. 2016), we imporved the previous optogenetic AKT system (Ong et al. 2016) with two tandom CIBNs (referred to as optoAKT in this work) (Supplemental Fig. S1A). Consistent with previous studies, the association of CIBN and CRY2 took about 1 second and the CIBN-CRY2 complex dissociated in the dark within 10 minutes (Kennedy et al. 2010; Zhang et al. 2014). The fusion of Raf or AKT does not affect the association and dissociation kinetics of CIBN and CRY2 and multiple cycles of CRY2-CIBN association and dissociation can be triggered by alternating light-dark treatment (Supplemental Fig. S1B-S1D, Movie S1, S3). Activation of optoRaf and optoAKT resulted in nuclear translocation of ERK-EGFP (Fig. 1A, Movie S2) and nuclear export of FOXO3-EGFP (Fig. 1B, Movie S4) resolved by live-cell fluorescence imaging, indicative of activation of the ERK and AKT signaling pathways, respectively.

**Figure 1.**
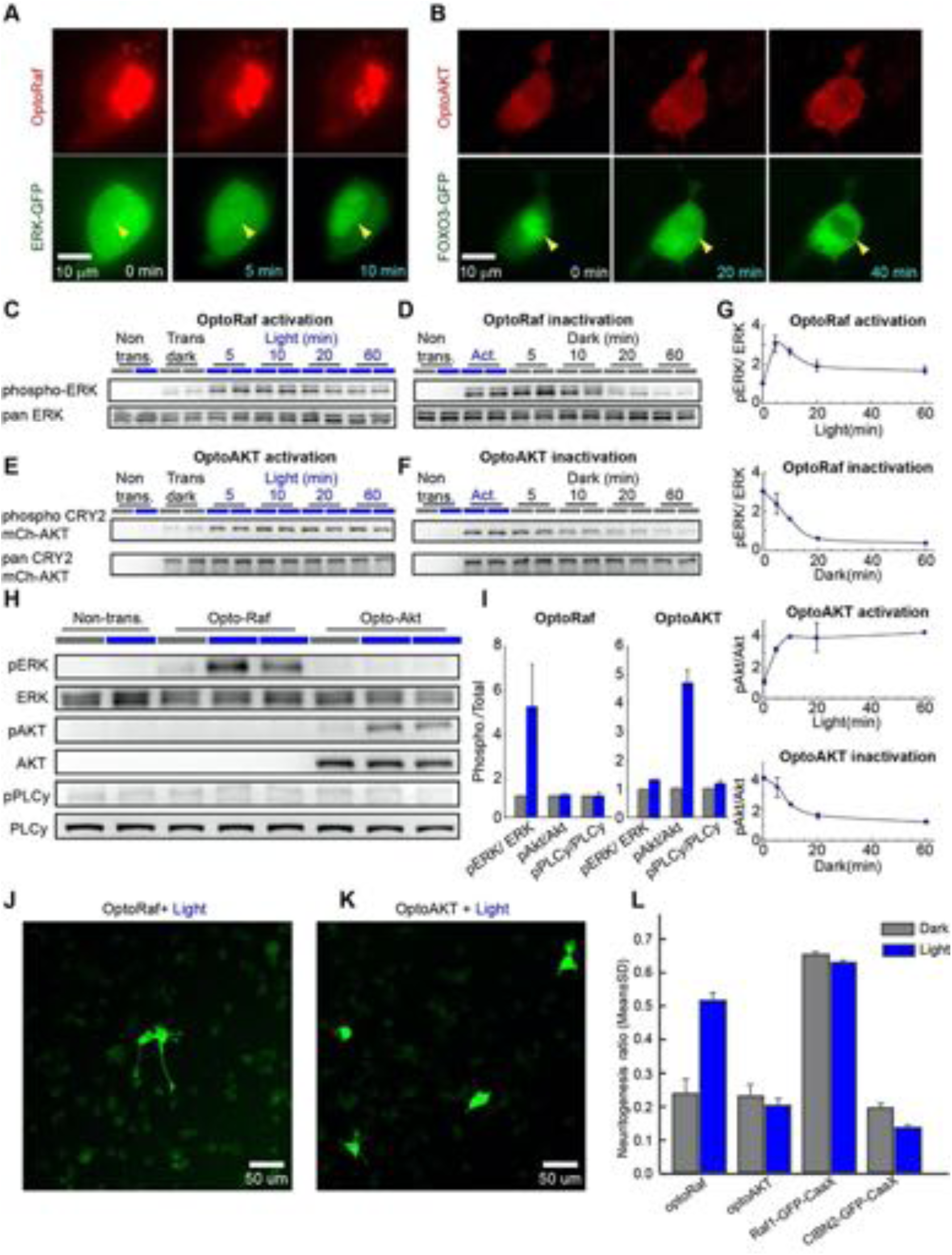
OptoRaf and optoAKT specifically activate the ERK and AKT subcircuits, respectively. (*A*) Activation of optoRaf benchmarked with ERK-GFP nuclear translocation. (*B*) Activation of optoAKT benchmarked with FOXO3-EGFP nuclear export. Scale bars = 10 µm. (*C*) Western blot analysis of the pERK and ERK activities in response to time-stamped activation of optoRaf. Blue light (0.5 mW/cm^2^) was applied for 5, 10, 20, and 60 min to HEK293T cells transfected with optoRaf. Non-transfected cells or optoRaf-transfected cells (dark) were used as negative controls. (*D*) Inactivation of the pERK activity after blue light was shut off. (*E*) Western blot analysis of the pAKT and AKT activities in response to time-stamped activation of optoAKT. Cells were treated with identical illumination scheme in (C). (*F*) Inactivation of the pAKT activity after blue light was shut off. (*G*) Quantification of the pERK activity (top two panels) and pAKT (bottom two panels) upon optoRaf and optoAKT activation, respectively. Both optoRaf and optoAKT show rapid (less than 5 min) and reversible activation patterns (*N* = 2). (*H*) OptoRaf and optoAKT do not show cross activity at the level of ERK and AKT. Neither optoRaf nor optoAKT can cause PLCy phosphorylation. Cells were exposed to blue light (0.5 mW/cm^2^) for 10 min before lysis. (*I*) Quantification of the phosphorylated protein level in (H) (*N* = 2). (*J, K*) PC12 cells transfected with either optoRaf (J) or optoAKT (K) were treated by blue light for 24 h (0.2 mW/cm^2^). Scale bars = 50 µm. (*I*) Quantification of the neuritogenesis ratio of PC12 cells transfected with optoRaf or optoAKT. A membrane-targeted Raf (Raf1-GFP-CaaX) causes constitutive neuritogenesis independent of light treatment, whereas the no-Raf (CIBN2-GFP-CaaX) control does not increase the neuritogenesis ratio under light or dark treatment.

Western blot analysis on pERK (activated by optoRaf) in HEK293 cells showed that pERK activity (Fig. 1C) increased within 10 min blue light stimulation and returned to the basal level 30 min after the blue light was shut off (Fig. 1D). There was a slight decrease of pERK activity upon optoRaf activation for over 10 min, likely due to a negative feedback, which has been consistently observed in previous studies (Zhou et al. 2017). On the other hand, continuous light illumination maintained a sustained activation of pCRY2-mCh-AKT (referred to as optoAKT in this work) within an onset of 10 min (Fig. 1E). The inactivation kinetics of pAKT was 30 min, similar to that of pERK (Fig. 1F, 1G). Note we use only the phosphorylated and total forms of CRY2-mCh-AKT to quantify the light response of optoAKT because the endogenous AKT does not respond to light.

### optoRaf and optoAKT do not show crosstalk activity at the pERK and pAKT level

Binding of neurotrophins to their receptor activates multiple downstream signaling subcircuits including the Raf/MEK/ERK, AKT, and phospholipase Cγ (PLCγ) pathways. Delineation of signaling outcomes of individual subcircuits remains difficult with pharmacological assays given the unpredictable off-targets of small-molecule drugs. We hypothesized that optoRaf and optoAKT could delineate signaling outcomes because they bypass ligand binding and activate the intracellular signaling pathway. To test this hypothesis, we probed phosphorylated proteins including pERK, pAKT, and pPLCγ with WB analysis in response to light-mediated activation of optoRaf and optoAKT. Results show that optoRaf activation does not increase pAKT and pPLCγ (Fig. 1H, 1I). Similarly, optoAKT activation does not increase pERK or pPLCγ (Fig. 1H, 1I). Thus, at the ERK and AKT level, optoRaf and optoAKT do not show crosstalk activity in mammalian cells.

### Activation of optoRaf but not optoAKT enhances PC12 cell neuritogenesis

We verified that activation of optoRaf enhances PC12 cell neuritogenesis, which is consistent with previous studies (Zhang et al. 2014; Krishnamurthy et al. 2016). The neuritogenesis ratio is defined as the ratio between the number of transfected cells with at least one neurite longer than the size of the cell body and the total number of transfected cells. Twenty-four hours of blue light stimulation (0.2 mW/cm^2^) increased the neuritogenesis ratio from the basal level (0.24 ± 0.04) to 0.52 ± 0.03 (Fig. 1J, 1L). Light-mediated activation of optoAKT, on the other hand, did not increase the neuritogenesis ratio (0.23 ± 0.04 in the dark versus 0.20 ± 0.02 under light) (Fig. 1K, 1L). A membrane-targeted Raf1 (Raf1-GFP-CaaX) was used as a positive control, which caused significant neurite outgrowth independent of light treatment (0.65 ± 0.01 in the dark versus 0.63 ± 0.01 under light). Expression of CIBN2-GFP-CaaX (without CRY2-Raf1), a negative control, did not increase PC12 neurite outgrowth either in the dark (0.20 ± 0.02) or under light (0.14 ± 0.01) (Fig. 1L).

### Activation of optoRaf but not optoAKT increases sensory neuron dendrite branching in fly larvae

To determine the efficacy of the optogenetic tools *in vivo*, we generated transgenic flies with inducible expression of optoRaf (*UAS-optoRaf*) and optoAKT (*UAS-optoAKT*). We induced the expression of the transgenes in a type of fly sensory neurons, the dendritic arborization (da) neurons, which have been used extensively to study dendrite morphogenesis and remolding (Gao et al. 1999; Grueber et al. 2002; Sugimura et al. 2003; Kuo et al. 2005; Williams and Truman 2005; Kuo et al. 2006; Williams et al. 2006; Parrish et al. 2007). Using the *pickpocket (ppk)-Gal4*, we specifically expressed optoRaf in the class IV da (C4da) neurons, to test whether light stimulation would activate the Raf/MEK/ERK pathway. At 72 hours after egg laying (h AEL), wild-type (WT) and optoRaf-expressing larvae were anesthetized with ether and subjected to wholefield continuous blue light for 5 min, while as a control, another transgenic group was incubated in the dark. The larval body walls were then dissected and immunostained with the pERK1/2 antibody, as a readout of the Raf/MEK/ERK pathway activation. We found that, compared with the optoRaf-expressing larvae incubated in the dark and WT larvae, light stimulation significantly increased the pERK signal in the cell body of C4da neurons in optoRaf-expressing larvae (Fig. 2A), leading to 3.5-fold increase in fluorescence intensity (Fig. 2B). Similarly, in C4da neurons expressing optoAKT, the 5-min blue light stimulation significantly increased the fluorescence intensity of phospho-p70 S6 kinase (phospho-p70^S6K^) (Fig. 2C, 2D), which functions downstream of AKT (Lizcano et al. 2003; Miron et al. 2003). These results collectively demonstrate that blue light is sufficient to activate the optogenetic effectors in flies *in vivo*.

**Figure 2.**
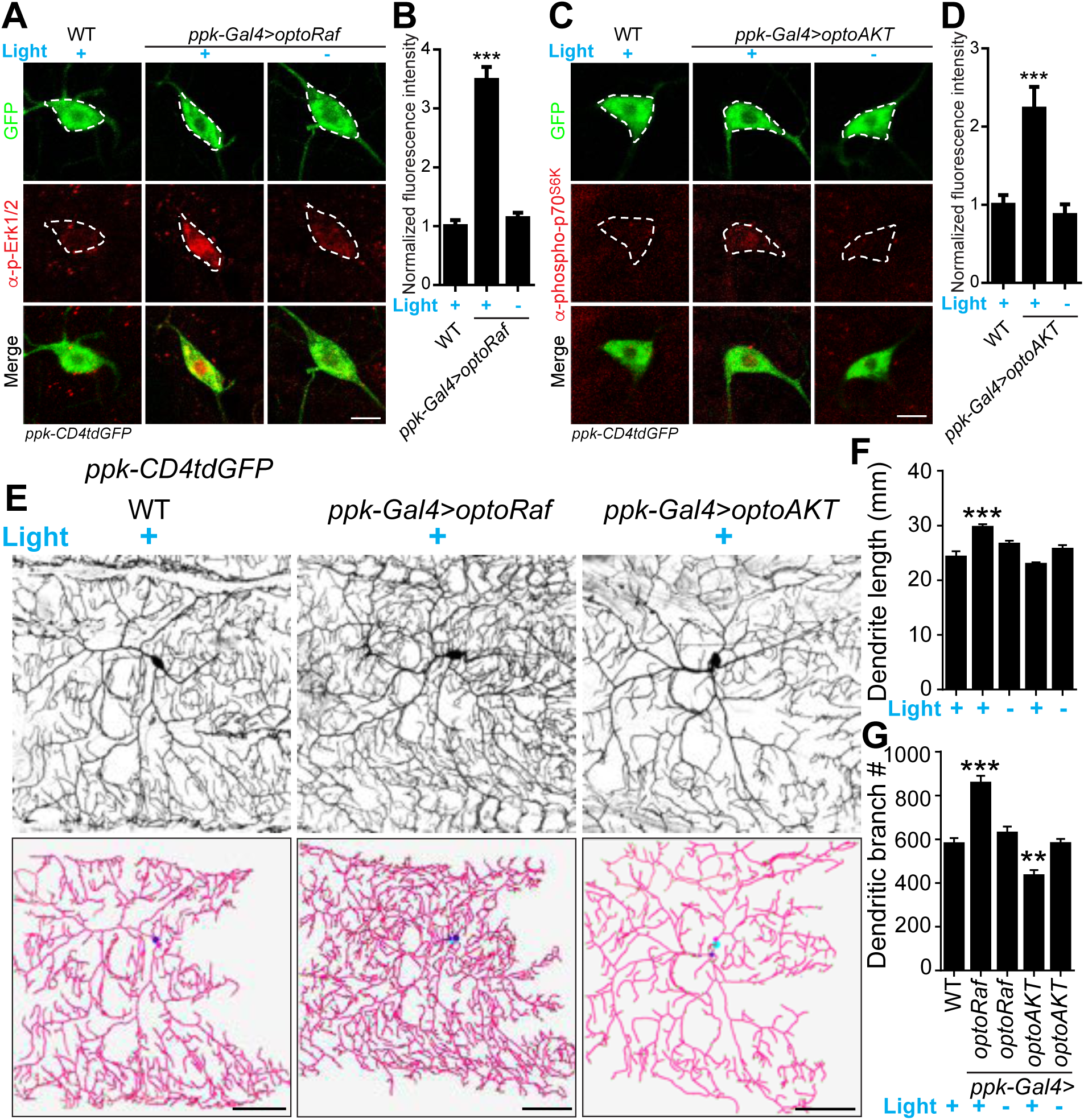
Activation of optoRaf but not optoAKT increases C4da neuron dendrite complexity. (*A, B*) Blue light stimulation activates optoRaf in flies *in vivo*. (A) The body walls from WT and optoRaf expressing larvae were dissected and stained for pERK1/2. After 5-min continuous light stimulation, the relative intensity of pERK is significantly increased in the optoRaf-expressing C4da neurons (labeled by *ppk-CD4tdGFP*). C4da neuron cell bodies are outlined by dashed white lines. Scale Bar = 10 µm. (B) Qualification of pERK fluorescence intensity in (A). The intensity of pERK in optoRaf expressing larvae was normalized to that of WT. *N* = 18 neurons. (*C, D*) The 5-min light stimulation is sufficient to activate optoAKT *in vivo*. (C) The larval body walls from WT and optoAKT expressing larvae were dissected and stained for phospho-p70^S6K^, a downstream component of the AKT pathway. Light stimulation increases phospho-p70^S6K^ signal intensity in C4da neurons expressing optoAKT. C4da neuron cell bodies are outlined by dashed white lines. Scale Bar = 10 µm. (D) Qualification of phospho-p70^S6K^ fluorescence intensity in (C). The intensity of phospho-p70^S6K^ in optoAKT expressing larvae was normalized to that of WT. WT (light) *N* = 17, *optoAKT* (light) *N* = 19, *optoAKT* (dark) *N* = 18 neurons. (*E*-*G*) Activation of Raf/MEK/ERK but not AKT signaling by 72 hours’ light stimulation increases dendrite outgrowth and branching in C4da neurons. (E) Representative images of C4da neurons from WT, optoRaf and optoAKT expressing larvae with 72 hours’ light stimulation and the unstimulated controls. Neurons were reconstructed with Neurostudio. Scale bar = 50 µm. (F) Quantification of total dendrite length of C4da neurons. (G) Qualification of dendritic branch number. WT (light) *N* = 21, *optoRaf* (light) *N* = 20, *optoRaf* (dark) *N* = 21, *optoAKT* (light) *N* = 21, *optoAKT* (dark) *N* = 20 neurons. All data are mean ± SEM. The data were analyzed by one-way ANOVA followed by Dunnett’s multiple comparisons test, \*\**P* < 0.01, \*\*\**P* < 0.001.

We next investigated if optoRaf or optoAKT activation would affect neural development such as dendrite morphogenesis. We labeled C4da neurons with *ppk-CD4tdGFP* and reconstructed the dendrites of the lateral C4da neurons – v’ada. Without light stimulation, the dendrite complexity of neurons in transgenic larvae was comparable to that of WT (Fig. 2F, 2G). However, optoRaf activation resulted in a significant increase in both total dendrite length and branch number, while optoAKT activation exhibited a slight reduction in dendritic branching (Fig. 2E-2G). These results confirm the possibility of independently activating the Raf/MEK/ERK and AKT signaling pathways in flies with our optogenetic tools, prompting us to test the feasibility of their *in vivo* applications, such as promoting axon regeneration with high spatial and temporal resolution.

### Activation of optoRaf or optoAKT results in enhanced axon regeneration in the PNS

Administration of neurotrophins to damaged peripheral neurons results in functional regeneration of sensory axons into the adult spinal cord in rat (Ramer et al. 2000). Here, our photoactivatable transgenic flies empower precise spatiotemporal control of the neurotrophic signaling in live animals. To test whether light-mediated activation of the Raf/MEK/ERK or AKT signaling subcircuits would also promote axon regrowth, we used a previously described *Drosophila* da sensory neuron injury model (Song et al. 2012; Song et al. 2015). Da neurons have been shown to possess distinct regeneration capabilities among different sub-cell types, and between the PNS and CNS, resembling mammalian neurons (Song et al. 2012; Song et al. 2015). In particular, the C4da neurons regenerate their axons robustly after peripheral injury, while the C3da neurons largely fail to regrow. Moreover, the axon regeneration potential of C4da neurons is also diminished after CNS injury. First, we asked whether optoRaf or optoAKT activation can enhance axon regeneration in the regeneration-competent C4da neurons in the PNS. We severed the axons of C4da neurons (labeled with *ppk-CD4tdGFP*) with a two-photo laser at 72 h AEL, verified axon degeneration at 24 h after injury (AI) and assessed axon regeneration at 48 h AI. At this time point, about 79% C4da neurons in WT showed obvious axon regrowth, and the regeneration index (Song et al. 2012; Song et al. 2015), which refers to the increase in axon length normalized to larval growth (Supplementary Fig. 2A, 2B, and Materials and Methods), was 0.381 ± 0.066 (Fig. 3A-3C). Strikingly, C4da neurons expressing optoRaf or optoAKT showed further enhanced regeneration potential in response to blue light, leading to a significant increase in the regeneration index (*optoRaf*: 0.682 ± 0.115; *optoAKT*: 0.735 ± 0.078), while there was no difference between WT and unstimulated transgenic flies (Fig. 3A-3C). In order to test the potential synergy between optoRaf and optoAKT, we co-expressed both in C4da neurons. While there is a slight increase in the regeneration percentage, activation of both ERK and AKT pathways in the same neuron did not further increase the regeneration index (0.7387 ± 0.08390) (Fig. 3A-3C). This suggests that these two subcircuits may share the same downstream components in promoting axon regeneration. Alternatively, activation of the ERK or AKT pathway by optogenetics may be strong enough to cause a saturation effect in C4da neurons axon regeneration.

**Figure 3.**
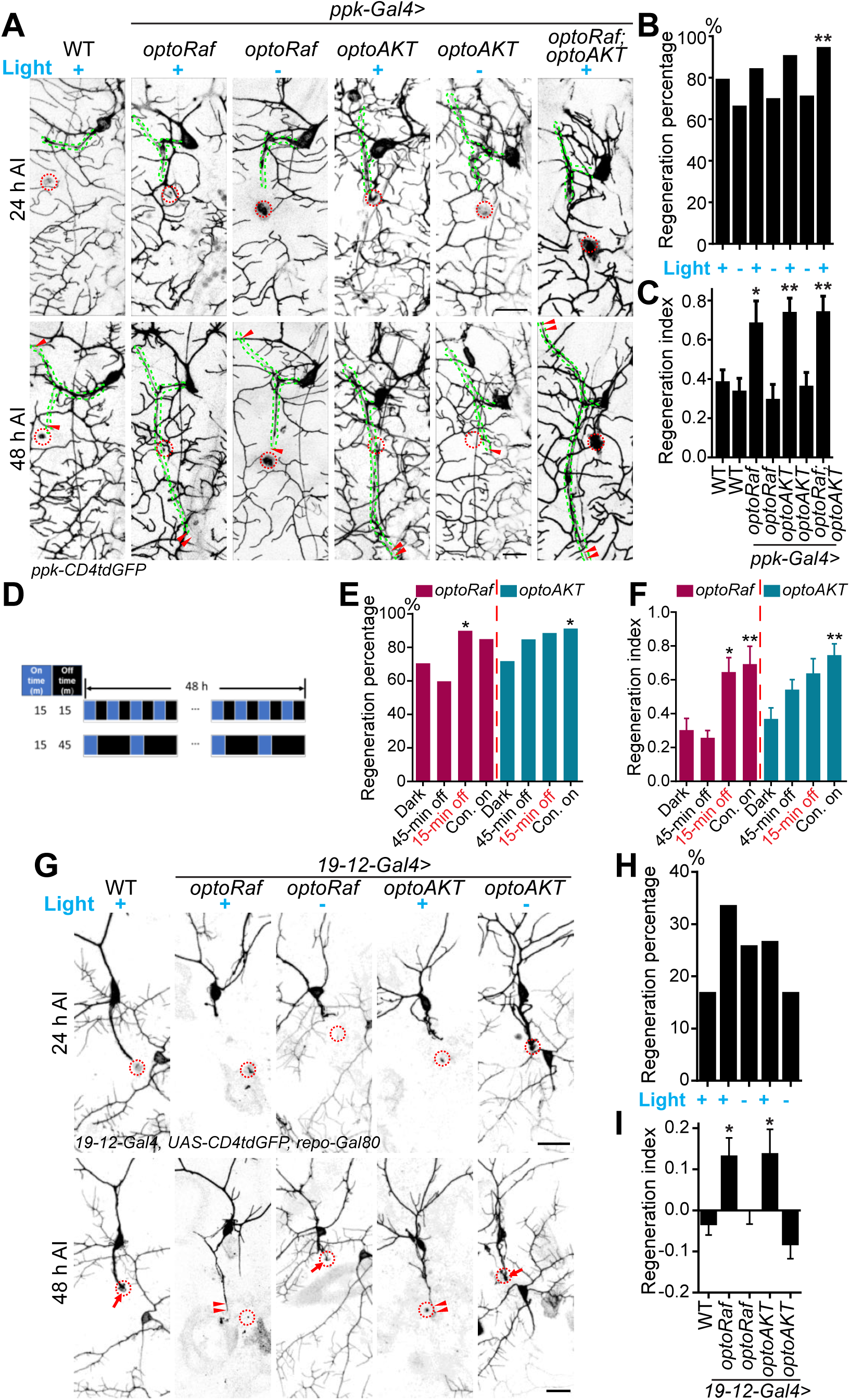
Light-stimulated optoRaf or optoAKT enhances axon regeneration in the PNS. (*A*-*C*) Compared to WT, C4da neurons expressing optoRaf or optoAKT show significantly increased axon regeneration in response to blue light. No enhancement was observed in the unstimulated controls. (A) C4da neuron axons were severed and their regeneration was assayed at 48 h AI. The injury site is marked by the dashed circle and regenerating axons are marked by arrowheads. Axons are outlined with dashed green lines. Scale bar = 20 µm. (B) The regeneration percentage of light-stimulated transgenic groups is not significantly higher than WT. Fisher’s exact test, *P* = 0.2199, *P* = 0.6026, *P* = 0.8829, *P* = 0.1445, *P* = 0.7886, *P* = 0.0025. (C) Qualification of C4da neuron axon regeneration by the regeneration index. WT (light) *N* = 33, WT (dark) *N* = 41, *optoRaf* (light) *N* = 31, *optoRaf* (dark) *N* = 36, *optoAKT* (light) *N* = 52, *optoAKT* (dark) *N* = 41, *optoRaf + optoAKT* (light) *N*=51 neurons. Data are mean ± SEM, analyzed by one-way ANOVA followed by Dunnett’s multiple comparisons test. (*D*-*F*) After injury, larvae were subjected to programmed light and dark cycles for a total of 48 h. The intermittent light stimulation promotes axon regrowth in optoRaf expressing larvae similar to constant light when the off-time is 15 min. (D) The intermittent patterns of the light stimulus. (E) Compared to larvae incubated in dark, light stimulation is capable of increasing the percentage of regenerated axons. Fisher’s exact test, *P* = 0.3357, *P* = 0.0422, *P* = 0.1673; *P* = 0.1414, *P* = 0.0639, *P* = 0.0307. (F) Qualification of C4da axon regeneration by the regeneration index. *OptoRaf* (dark) *N* = 36, *optoRaf* (45-min off) *N* = 43, *optoRaf* (15-min off) *N* = 36, *optoRaf* (Con. on) *N* = 31, *optoAKT* (dark) *N* = 41, *optoAKT* (45-min off) *N* = 49, *optoAKT* (15-min off) *N* = 40, *optoAKT* (Con. on) *N* = 52 neurons. Data are mean ± SEM, analyzed by one-way ANOVA followed by Dunnett’s multiple comparisons test. (*G*-*I*) Blue light stimulation significantly enhances axon regeneration in the regeneration-incompetent C3da neurons. (G) C3da neuron axon degeneration was verified at 24 h AI and axon regeneration was assessed at 48 h AI. The injury site is marked by the dashed circle, regenerated axons are demarcated by arrowheads, and arrow marks non-regenerated axons. Scale bar = 20 µm. (H) The regeneration percentage is not significantly different. Fisher’s exact test, *P* = 0.0874, *P* = 0.9910, *P* = 0.2972, *P* = 1.0000. (I) Qualification of axon regeneration by the regeneration index. WT (light) *N* = 41, *optoRaf* (light) *N* = 36, *optoRaf* (dark) *N* = 39, *optoAKT* (light) *N* = 34, *optoAKT* (dark) *N* = 38 neurons. Data are mean ± SEM, analyzed by one-way ANOVA followed by Dunnett’s multiple comparisons test. \**P* < 0.05, \*\**P* < 0.01. See also Supplementary Fig. 2.

The light stimulation paradigm used in the aforementioned *in vivo* experiments was constant blue light applied immediately after injury. We reason that intermittent light stimulation may provide insights into the signaling kinetics *in vivo* and fine-tune axon regeneration dynamics. Therefore, instead of constant blue light on, we delivered two sets of programmed light patterns to injured larvae, 15 min on-15 min off or 15 min on-45 min off per cycle for 48 h (Fig. 3D). We found that, for optoRaf-expressing C4da neurons, when the off-time was 15 min, the intermittent light stimulation was sufficient to accelerate axon regrowth, with both the regeneration index (0.6352 ± 0.09627) and regeneration percentage significantly increased compared to larvae incubated in the dark (Fig. 3E, 3F). However, when the off-time was 45 min, the intermittent light failed to promote axon regeneration (Fig. 3E, 3F), suggesting a threshold effect. On the other hand, C4da neurons expressing optoAKT displayed a graded response: a moderate increase of regeneration index (0.6278 ± 0.09801) in response to the 15 min on-15 min off light and a smaller uptick to the 15 min on-45 min off light; both were less effective than the constant light stimulation (Fig. 3E, 3F). These results suggest that although higher frequency of light stimulation generally resulted in stronger regeneration potential in the transgenic flies, constant light was not always required for maximum axon regeneration. Moreover, optoRaf and optoAKT differ in their signaling kinetics during regeneration, showing a gated versus graded response, respectively.

We next determined whether optoRaf or optoAKT activation would trigger regeneration in C3da neurons, which are normally incapable of regrowth (Song et al. 2012). C3da neurons were labeled with *19-12-Gal4, UAS-CD4tdGFP, repo-Gal80* and injured using the same paradigm as C4da neurons. Compared to WT, which exhibited poor axon regeneration ability demonstrated by the low regeneration percentage and the negative regeneration index (−0.03201 ± 0.02752) (Fig. 3G-3I), light stimulation significantly increased the regeneration index in optoRaf- or optoAKT-expressing larvae to 0.1298 ± 0.04637 or 0.1354 ± 0.06161, respectively (Fig. 3G-3I). These results indicate that optoRaf and optoAKT activation not only accelerates axon regeneration but also converts regeneration cell-type specificity.

### Spatial activation of optoRaf or optoAKT improves pathfinding of regenerating axons

While C4da neurons are known to possess the regenerative potential, it is unclear whether the regenerating axons navigate correctly. To address this question, we focused on v’ada – the lateral C4da neurons. Uninjured v’ada axons grow ventrally, showing a typical turn and then join the axon bundle with the ventral C4da neurons (Supplementary Fig. 2A). We found that their regenerating axons preferentially regrew away from the original ventral trajectory. More than 60% v’ada axons bifurcated and formed two branches targeting opposite directions (Fig. 4A, 4B white bar). In the majority cases in WT, the ventral branch, which extends towards the correct trajectory, regenerated less frequently than the dorsal branch, with 15% v’ada containing only the ventral branch (Fig. 4A, 4B black bar). One possibility is that the ventral branch encounters the injury site, which may retard its elongation. As a result, only a minority of regenerating axons are capable of finding the correct path. The poor pathfinding of regenerating axons was similar among WT and the transgenic larvae, regardless of whether incubated with whole-field light or in the dark (Fig. 4B). Thus, proper guidance of the regenerating axons towards the correct trajectory remained to be resolved.

**Figure 4.**
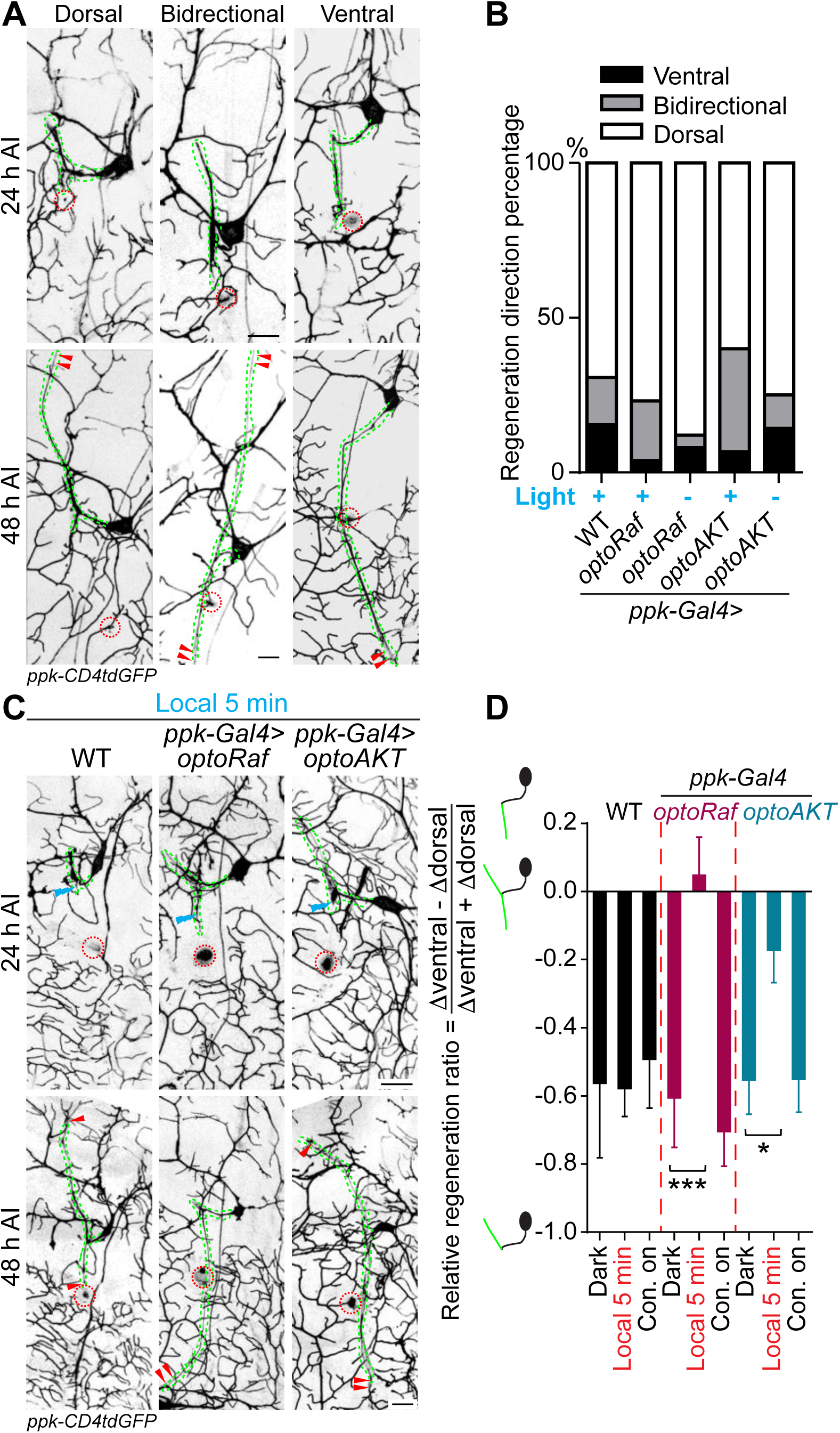
Local optogenetic stimulation conveys guidance instructions to regenerating axons. (*A, B*) Regenerating axons prefer to regrow away from the original trajectory, with only a minority of axons finding the correct path. (A) Representative images of axons retracting or bifurcating at 24 h AI. At 48 h AI in WT, regenerating axons extend dorsally, ventrally, or both directions. The injury site is marked by the dashed circles and regenerating axons are marked by arrowheads. Axons are outlined with dashed green lines. Scale bar = 20 µm. (B) Light stimulation fails to increase the percentage of axons regrowing towards the right direction. The percentage of axons extending towards the correct trajectory (ventral + both) were analyzed by Fisher’s exact test, *P* = 0.5318, *P* = 0.1033, *P* = 0.7778, *P* = 0.6363. WT (light) *N* = 26, *optoRaf* (light) *N* = 26, *optoRaf* (dark) *N* = 25, *optoAKT* (light) *N* = 45, *optoAKT* (dark) *N* = 28 neurons. (*C, D*) Restricted local activation of optoRaf or optoAKT significantly increases the relative regeneration ratio. The ratio is defined to weigh the regeneration potential of the ventral branch against the dorsal branch. (C) A single pulse of light stimulation delivered specifically on the ventral axon branch at 24 h AI (blue flash symbol) is capable of promoting preferential extension of regenerating axons in optoRaf or optoAKT expressing larvae. (D) Qualification of the relative regeneration ratio of v’ada. WT (dark) *N*= 32, WT (local 5 min) *N* = 32, WT (Con. on) *N* = 33, *optoRaf* (dark) *N* = 33, *optoRaf* (local 5 min) *N* = 35, *optoRaf* (Con. on) *N* = 35, *optoAKT* (dark) *N* = 33, *optoAKT* (local 5 min) *N* = 34, *optoAKT* (Con. on) N = 36 neurons. Data are mean ± SEM, analyzed by one-way ANOVA followed by Dunnett’s multiple comparisons test, \**P* < 0.05, \*\*\**P* < 0.001.

We thus investigated whether spatially restricted activation of the neurotrophic signaling using our optogenetic system could guide the regenerating axons. To specifically enhance the regrowth of the ventral branch, we used a confocal microscope to focus the blue light (delivered by the 488 nm argon-ion laser) on the ventral branch for 5 min at 24 h AI. The lengths of both the ventral and dorsal branches were measured at 24 h AI and 48 h AI. We subtracted the increased dorsal branch length (Δdorsal) from the increased ventral branch length (Δventral), then divided that by the total increased length of these two branches (Fig. 4D). This value was defined as the relative regeneration ratio. If the dorsal branch exhibited more regenerative potential, the ratio would be negative; otherwise, it would be positive. Without light stimulation, the relative regeneration ratio of the transgenic larvae (*optoRaf*: −0.6062 ± 0.1453; *optoAKT*: - 0.5530 ± 0.1011) was comparable to that of WT (−0.5786 ± 0.08229) (Fig. 4C, 4D), confirming preferred regrowth of the dorsal branch. Strikingly, the 5-min local blue light stimulation significantly increased the ratio in optoRaf- or optoAKT-expressing v’ada (*optoRaf*: 0.04762 ± 0.1123; *optoAK*T: −0.1725 ± 0.09560), while this transient stimulation resulted in no difference in WT (−0.6018 ± 0.1290) (Fig. 4C, 4D). This result indicates that a single pulse of local light stimulation was sufficient to lead to preferential regrowth of the ventral branch. Notably, although whole-field light illumination could significantly promote axon regrowth, it failed to increase the relative regeneration ratio in transgenic larvae (Con.on *optoRaf*: −0.7048 ± 0.1015; Con. on *optoAKT*: −0.5517 ± 0.09644) (Fig. 4D), revealing the difference between activating the neurotrophic signaling in a whole neuron and a single lesioned axon branch. On the other hand, while a 5 min local light stimulation did not lead to an overall enhancement of axon regrowth, it provided adequate guidance instructions for the regenerating axons to make the correct choice.

### Activation of optoRaf or optoAKT promotes axon regeneration and functional recovery in the CNS

Achieving functional axon regeneration after CNS injury remains a major challenge in neural repair research. Motivated by the capacity of optoRaf and optoAKT to accelerate axon regeneration in the PNS, we went on to determine whether they also show efficacy after CNS injury. We focused on the axons of C4da neurons, which project into the ventral nerve cord (VNC) and form a ladder-like structure. Each pair of axon bundles correspond to one body segment in an anterior-posterior pattern (Fig. 5A) (Li et al., submitted). We injured the abdominal A6 and A3 bundles by laser as previously described (Song et al. 2012) (Li et al., submitted) (Supplementary Fig. 3), and confirmed axon degeneration at 24 h AI (Fig. 5A). At 48 h AI, we found that axons began to extend from the retracted axon stem and towards the commissure region. We defined a commissure segment as regenerated only when at least one axon extended beyond the midline of the commissure region or joined into other intact bundles (Supplementary Fig. 3). In WT, only 16% of lesioned commissure segments displayed obvious signs of regrowth (Fig. 5A, 5B). To quantify the extent of regrowth, we measured the length of the regrown axons and normalized that to the length of a commissure segment – regeneration index (Supplementary Fig. 3, Materials and Methods). After light stimulation, the regeneration indexes of the two transgenic lines (*optoRaf*: 5.375 ± 0.3391; *optoAKT*: 4.765 ± 0.4236) were significantly increased compared with the WT control (2.643 ± 0.3050), and the percentage of regenerating commissure segments also exhibited a mild increase in both optoRaf- and optoAKT-expressing larvae (Fig. 5A-5C). On the other hand, there was no significant difference between WT and the unstimulated transgenic flies (Fig. 5A-5C). This result suggests that both signaling subcircuits reinforce C4da neuron axon regeneration in the CNS.

**Figure 5.**
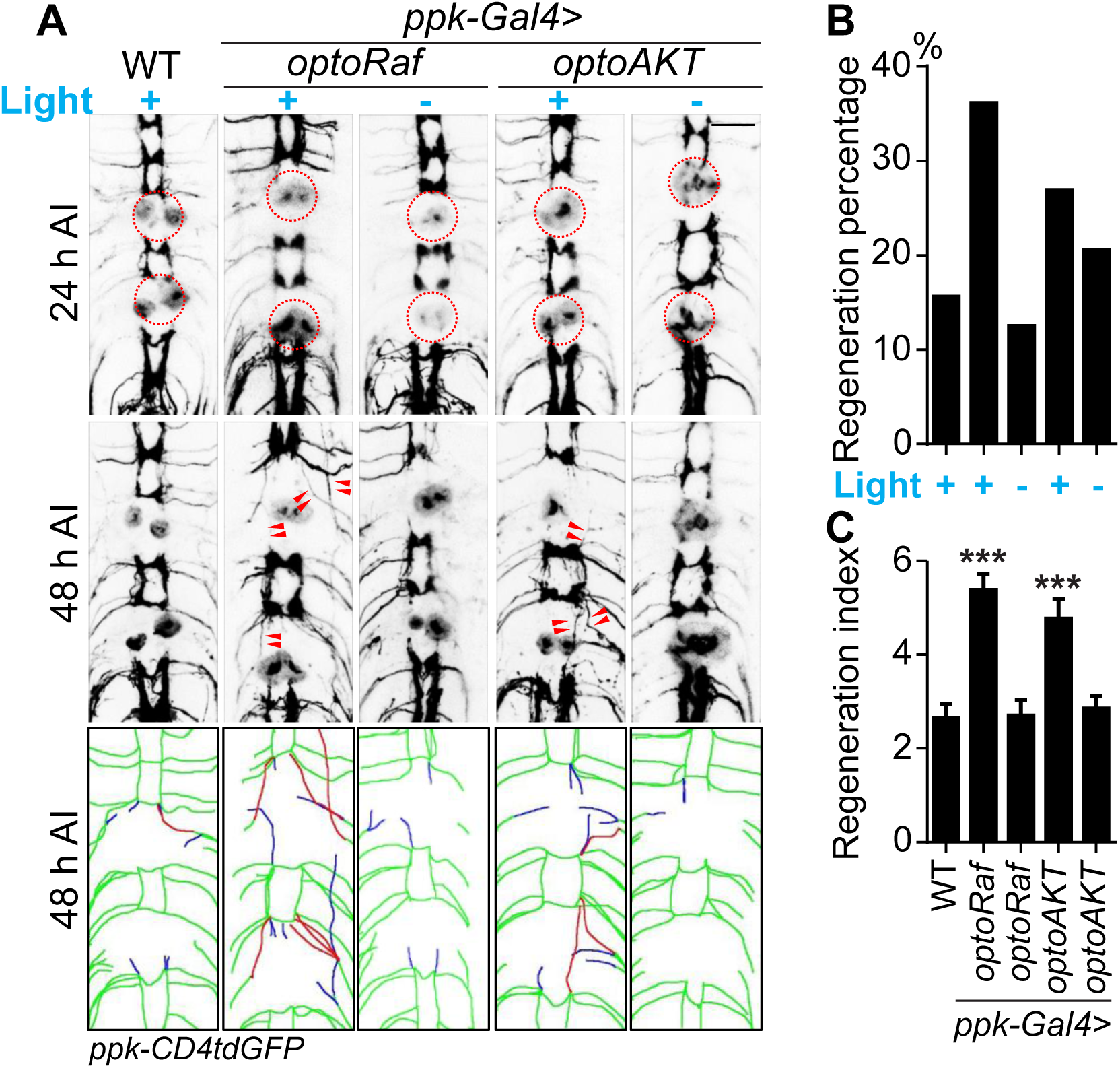
Activation of optoRaf or optoAKT promotes axon regeneration in the CNS. (*A*-*C*) Light stimulation significantly enhances axon regeneration in the VNC of optoRaf or optoAKT expressing larvae. (A) Complete degeneration in A3 and A6 commissure segments was confirmed at 24 h AI and regeneration of these two segments was assayed independently at 48 h AI. The injury site is marked by the dashed circle and regenerating axons are labeled by arrowheads. In the schematic diagrams, regrowing axons that reached other bundles and thus define a regenerating commissure segment are highlighted in red, while other regrowing axons are illustrated in blue. Scale bar = 20 µm. (B) The regeneration percentage is not significantly different. Fisher’s exact test, *P* = 0.0560, *P* = 0.7192, *P* = 0.2908, *P* = 0.6013. (C) Qualification of axon regeneration in VNC by the regeneration index. WT (light) *N* = 32, *optoRaf* (light) *N* = 36, *optoRaf* (dark) *N* = 32, *optoAKT* (light) *N* = 26, *optoAKT* (dark) *N* = 34 segments. Data are mean ± SEM, analyzed by one-way ANOVA followed by Dunnett’s multiple comparisons test, \*\*\**P* < 0.001. See also Supplementary Fig. 3.

We then tested whether the axon regrowth in the CNS induced by optoRaf or optoAKT activation leads to behavioral improvement. We utilized a recently established behavioral recovery paradigm based on larval thermonociception (Fig. 6A, Materials and Methods) (Li et al., submitted). In brief, we injured the A7 and A8 C4da neuron axon bundles in the VNC, which correspond to the A7 and A8 body segments in the periphery. We then assessed the nociceptive behavior in these larvae in response to a 47 °C heat probe applied at the A7 or A8 segments at 24 and 48 h AI. Since C4da neurons are essential for thermonociception, injuring A7 and A8 axon bundles in the VNC would lead to an impaired nociceptive response to the heat probe specifically at body segments A7 and A8. Indeed, all the injured larvae exhibited diminished response at 24 h AI, while the total score is approaching 3 in uninjured WT larvae (Fig. 6B). At 48 h AI, substantial recovery was observed in the two transgenic groups with light stimulation, whereas WT showed very limited response and a low recovery percentage (Fig. 6B, 6C). Both the response score and the percentage of larvae exhibiting behavioral recovery in these two groups were more than twice as that of the WT, while the unstimulated groups were comparable to WT. Altogether, these results demonstrate that our optogenetic system empowers ligand-free and non-invasive control of the Raf/MEK/ERK and AKT pathways in flies, which not only promote axon regeneration after injury but also benefit functional recovery, suggesting that the regenerated axons may rewire and form functional synapses.

**Figure 6.**
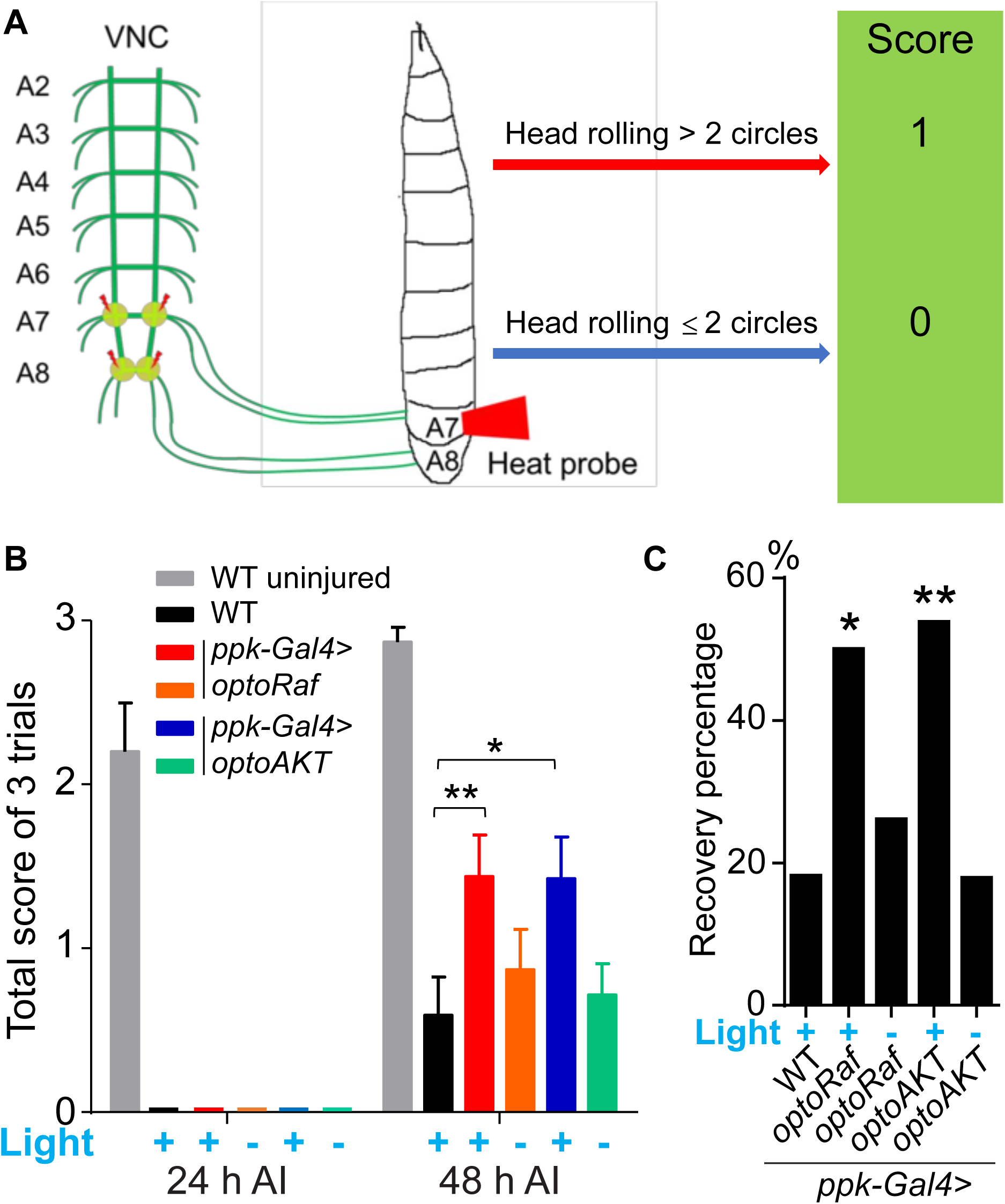
Activation of optoRaf or optoAKT promotes functional regeneration after CNS injury. (*A*) A schematic drawing of the behavioral recovery test. The A7 and A8 C4da neuron axon bundles (corresponding to the A7 and A8 body segments) in the VNC were injured by laser and the larva was then subjected to three consecutive trials at 24 and 48 h AI, respectively. In each trial, a 47°C heat probe was applied at the A7 or A8 segments. A fully recovered larva would produce a stereotypical rolling behavior in response to the heat probe and would be scored as “1”, otherwise as “0”. If the total score of the three trials was below 1 at 24 h AI but increased to 2 or 3 at 48 h AI, the larva was defined as recovered. (*B, C*) Behavioral recovery test was performed at 24 h and 48 h after VNC injury (A7 and A8 bundles). Larvae expressing optoRaf or optoAKT exhibit significantly accelerated recovery in response to light stimulation. (B) Qualification of the total scores at each time point. WT (uninjured) *N* = 15, WT (light) *N* = 22, *optoRaf* (light) *N* = 32, *optoRaf* (dark) *N* = 23, *optoAKT* (light) *N* = 26, *optoAKT* (dark) *N* = 28. Data are mean ± SEM, analyzed by two-way ANOVA followed by Tukey’s multiple comparisons test. (C) Qualification of the recovery percentage. The data were analyzed by Fisher’s exact test, *P* = 0.0174, *P* = 0.5237, *P* = 0.0052, *P* = 0.9763. \**P* < 0.05, \*\**P* < 0.01.

## Discussion

Neurotrophins are known to activate Trk receptors and trigger the Ras/MEK/ERK, AKT, and PLCy pathways which are involved in cell survival, neural differentiation, axon and dendrite growth and sensation (Bibel and Barde 2000; Huang and Reichardt 2001; Chao 2003; Cheng et al. 2011; Joo et al. 2014). Here, we used optogenetic systems to achieve specific and reversible activation of the neurotrophin subcircuits including the Raf/MEK/ERK (via optoRaf) and AKT (via optoAKT) signaling pathways. We further verified that optoRaf and optoAKT did not show crosstalk at the level of phosphorylated ERK and AKT proteins, and activation of optoRaf but not optoAKT promoted PC12 cell differentiation.

After spinal cord injury, the synthesis of neurotrophins is elevated to support axon regrowth (Cho et al. 1998; Hayashi et al. 2000; Fukuoka et al. 2001; Fang et al. 2017). AKT signaling, which functions downstream of Trk receptors, was reported to accelerate axon regeneration in fly and mammals (Song et al. 2012; Guo et al. 2016; Miao et al. 2016). However, the role of Raf/MEK/ERK signaling played during nerve repair is controversial. Although some studies revealed that ERK is involved in axon extension, others suggested that ERK activation impedes axon regeneration and functional recovery (Markus et al. 2002; Huang et al. 2017; Cervellini et al. 2018). To specifically evaluate the efficacy of Raf/MEK/ERK and AKT signaling in promoting axon regeneration, we generated fly strains with tissue-specific expression of optoRaf or optoAKT and found that light stimulation was sufficient to activate the corresponding downstream components in fly larvae *in vivo*. Consistent with previous studies (He and Jin 2016), we found that AKT activation resulted in significantly increased axon regeneration in C4da neurons as well as the regeneration-incompetent C3da neurons. Interestingly, we found that C4da and C3da neurons expressing optoRaf also exhibited greater regeneration potential in response to light stimulation. This result also corroborates with a previous finding that activated B-RAF signaling enables axon regeneration in the mammalian CNS (O’Donovan et al. 2014). We speculate that the differential outcomes of ERK activation on axon regeneration may be due to the different injury models used, and the strength and cell type origin of ERK signaling.

The regenerative capacity varies significantly among different neuronal subtypes, as well as the PNS and CNS. Although the administration of neurotrophins enhances axon regeneration in peripheral neurons, its capacity to promote functional regeneration in the CNS is limited, in part due to the inaccessibility of neurotrophins to reach injured axons (physical barrier) (Silver and Miller 2004; Yiu and He 2006) and innate inactivation of the regenerating program in CNS (Lu et al. 2014). OptoRaf and optoAKT could be used to address both issues by direct delivery of light (rather than ligand) to reactivate the regenerating program and thereby significantly increase neural regeneration in the CNS as well. We further showed that activation of the Raf/MEK/ERK or AKT subcircuit was capable of improving behavioral performance in fly larvae, suggesting that it may promote synapse regeneration leading to functional recovery.

Ineffective functional recovery at least partially results from the inappropriate pathfinding of the regenerative neurons. As shown in this study, the majority of regenerating C4da neuron axons preferentially grew away from their original trajectory. We surprisingly found that delivering a 5-min light stimulation to the ventral branch, which extended towards the correct direction, was sufficient to convey guidance instructions and increase the preferential elongation of the ventral branch against the dorsal branch. Correct guidance cannot be achieved by whole-body administration of pharmacological reagents. Similarly, when casting blue light on the whole transgenic larvae, light stimulation must be given at a high frequency to promote axon regrowth (there is a threshold for the light off-time), and the dorsal branch extension was also dominant in this case. This result highlights the importance and necessity of restricted activation of neurotrophic signaling. Indeed, the strength and location of Raf/MEK/ERK and AKT activation during axon regeneration may be important to the functional consequences. Notably, although the transient restricted stimulation likely affects the decision-making of the growth cone at the branching point, constant light is still required to increase overall axon regeneration.

Neurotrophins are engaged in a variety of important cellular processes, and their physiological concentration is essential for the normal function of both neurons and non-neuronal cells (Rose et al. 2003; Xiao et al. 2010; Poyhonen et al. 2019). Despite exhibiting substantial efficacy for enhancing nerve regeneration, neurotrophin-based therapeutic applications have been confronted with a number of obstacles such as their nociceptive effects and lack of strategy for localized signaling activation (Aloe et al. 2012; Mitre et al. 2017; Mahar and Cavalli 2018; Sung et al. 2019). OptoRaf and optoAKT aim to improve neurotrophin signaling outcomes by preferentially activating the neuroregenerative program and enabling spatiotemporal control. Our systems offer insights into the ERK and AKT subcircuits and delineate their differential roles downstream of neurotrophin activation, as evidenced by the distinct functional outcomes of Raf/MEK/ERK and AKT signaling in several aspects. First, ERK signaling promoted PC12 cell neuritogenesis, which was not induced by AKT activation. Second, elevated ERK activity significantly increased dendritic complexity, while on the contrary, AKT activation led to decreased dendrite branching. Third, after injury, C4da neurons expressing optoRaf and optoAKT responded differently to intermitted light stimulation, suggesting that their strength and activation duration is differentially gauged during axon regeneration. These collectively suggest that, since Raf could be activated by membrane translocation as well as dimerization, CRY2 oligomerization could further lead to a more potent Raf. This multimodal activation mechanism may render that a threshold of optoRaf can be reached so that a saturated ERK activation could be achieved. On the other hand, AKT activation does not depend on dimerization and may display a graded response. As a result, optoAKT activates the AKT pathway in a dose-dependent manner and may not recapitulate the maximan activation of AKT. This work provides a proof-of-principle to use optogenetics to accelerate and navigate axon regeneration in mammalian injury models. Besides spatiotemporal control of the neurotrophic signaling, optoRaf and optoAKT allow for finetuning of the signaling activity with programmed light pattern during axon regeneration. Follow-up studies are warranted to determine how Raf/MEK/ERK and AKT subcircuits are involved in each process of nerve repair, including lesioned axon degeneration, regenerating axon initiation and extension, and the formation of new synapses and remyelination in mammals. Understanding the machinery will, in turn, allow better utilization and development of the optogenetic systems. Although intact optogenetics in larger mammals is limited by the poor penetration depth of blue light (less than 1 mm), we are excited to witness the rapid progress in implantable, wireless µLED devices (Jeong et al. 2015; Park et al. 2015b) and the integration of optogenetics with long-wavelength responsive nanomaterials such as the upconversion nanoparticles (He et al. 2015; Wu et al. 2016; Chen et al. 2018), both of which would facilitate precise delivery of light stimulation.

## Materials and Methods

### Fly stocks

*19-12-Gal4* (Xiang et al. 2010), *reop-Gal80* (Awasaki et al. 2008), *ppk-CD4-tdGFP* (Han et al. 2011), and *ppk-Gal4* (Han et al. 2011) have been previously described. To generate the *UAS-optoRaf and UAS-optoAKT* stocks, the entire coding sequences were cloned into the pACU2 vector, and the constructs were then injected (Rainbow Transgenic Flies, Inc). Randomly selected male and female larvae were used. Analyses were not performed blind to the conditions of the experiments. The experimental procedures have been approved by the Institutional Biosafety Committee (IBC) at the Children’s Hospital of Philadelphia.

### Sensory axon lesion in *Drosophila*

Da neuron axon lesion and imaging in the PNS was performed in live fly larvae as previously described (Song et al. 2012; Stone et al. 2014; Song et al. 2015). VNC injury was performed as previously described (Song et al. 2012) (Li et al., submitted). In brief, A3 and A6 axon bundles in the VNC were ablated with a focused 930-nm two-photon laser and full degeneration around the commissure junction was confirmed 24 h AI. At 48 h AI, axon regeneration of these two commissure segments were assayed independently of each other (Supplementary Fig. 3).

### Quantitative analyses of sensory axon regeneration in flies

Quantification was performed as previously described (Song et al. 2012; Song et al. 2015). Briefly, for axon regeneration in the PNS, we used “regeneration percentage”, which depicts the percent of regenerating axons among all the axons that were lesioned; “regeneration index”, which was calculated as an increase of “axon length”/”distance between the cell body and the axon converging point (DCAC)” (Supplementary Fig. 2A, 2B). An axon was defined as regenerating only when it obviously regenerated beyond the retracted axon stem, and this was independently assessed of the other parameters. The regeneration parameters from various genotypes were compared to that of the WT if not noted otherwise, and only those with significant differences were labeled with the asterisks. For VNC injury, the increased length of each axon regrowing beyond the lesion sites was measured and added together. To calculate the regeneration index, the sum was then divided by the distance between A4 and A5 axon bundles (Supplementary Fig. 3). Regeneration percentage was assessed independently of the regeneration index. A commissure segment was defined as regenerated only when at least one regenerating axon passed the midline of the commissure region or joined into other intact bundles (Supplementary Fig. 3).

### Live imaging in flies

Live imaging was performed as described (Emoto et al. 2006; Parrish et al. 2007). Embryos were collected for 2-24 hours on yeasted grape juice agar plates and were aged at 25 °C or room temperature. At the appropriate time, a single larva was mounted in 90% glycerol under coverslips sealed with grease, imaged using a Zeiss LSM 880 microscope, and returned to grape juice agar plates between imaging sessions.

### Behavioral assay

The behavioral test was performed to detect functional recovery after VNC injury as described (Li et al., submitted). A7 and A8 C4da neuron axon bundles in the VNC, which correspond to the A7 and A8 body segments in the periphery, were injured with laser (Fig. 6A). Since C4da neurons are essential for thermonociception, such lesion results in impaired nociceptive response to noxious heat at body segments A7 and A8. We assessed larva nociceptive behavior in response to a 47 °C heat probe at 24 and 48 h AI. At each time point, the larva was subjected to three consecutive trials, separated by 15 seconds (s). In each trial, the heat probe was applied at the A7 and A8 body segments for 5 s. If the larva produced head rolling behavior for more than 2 cycles, it would be scored as “1”, otherwise “0” (Fig. 6A). The scores of the three trials were combined and the total score at 24 h AI was used to determine whether A7 and A8 bundles were successfully ablated. A larva was defined as recovered only when its total score was below 1 at 24 h AI but increased to 2 or 3 at 48 h AI. Those failed to exhibit such improvement at 48 h AI were defined as unrecovered. All the injured larvae exhibited normal nociceptive responses when the same heat probe was applied at the A4 or A5 body segment at 24 h AI.

### Immunohistochemistry

Third instar larvae or cultured neurons were fixed according to standard protocols. The following antibodies were used: rabbit anti-Phospho-p44/42 MAPK (Erk1/2) (Thr202/Tyr204) (4370, 1:100, Cell Signaling), rabbit anti-Phospho-*Drosophila* p70 S6 Kinase (Thr398) (9209S, 1:200, Cell Signaling) and fluorescence-conjugated secondary antibodies (1:1000, Jackson ImmunoResearch).

### Cell culture and transfection

HEK293T cells were cultured in DMEM medium supplemented with 10% FBS, and 1x Penicillin-Streptomycin solution (complete medium). Cultures were maintained in a standard humidified incubator at 37 °C with 5% CO_2_. For western blots, 800 ng of DNA were combined with 2.4 µL Turbofect in 80 µL of serum-free DMEM. The transfection mixtures were incubated at room temperature for 20 minutes prior to adding to cells cultured in 35 mm dishes with 2 mL complete medium. The transfection medium was replaced with 2 mL serum-free DMEM supplemented with 1× Penicillin-Streptomycin solution after 3 hours of transfection to starve cells overnight. PC12 cells were cultured in F12K medium supplemented with 15% horse serum, 2.5% FBS, and 1× Penicillin-Streptomycin solution. (For PC12 neuritogenesis assays, 2400 ng of DNA were combined with 7.2 mL of Turbofect in 240 mL of serum-free F12K. The transfection medium was replaced with 2 mL complete medium after 3 hours of transfection to recover cells overnight.). Twenty-four hours after recovery in high-serum F12K medium [15% horse serum + 2.5% fetal bovine serum (FBS)], the cell culture was exchanged to a low-serum medium (1.5% horse serum + 0.25% FBS) to minimize the base-level ERK activation induced by serum.

### Optogenetic stimulation for cell culture

For Western blot analysis, transfected and serum-starved cells were illuminated for different time using a home-built blue LED light box emitting at 0.5 mW/cm^2^. For PC12 cell neuritogenesis assay, PC12 cells were illuminated at 0.2 mW/cm^2^ for 24 h with the light box placed in the incubator.

### Optogenetic stimulation for fly

The whole optogenetics setup is modified from previous work (Kaneko et al. 2017). Larvae were grown in regular brown food at 25 °C in 12 h-12 h light-dark cycle. At 72 h AEL, early 3rd instar larvae were transferred from food, anesthetized with ether for axotomy. After recovery in regular grape juice agar plates, larvae were kept in the dark or under blue light stimulation thereafter. A 470 nm blue LED (LUXEON Rebel LED) was set over the grape-agar plate for stimulation. The LED was mounted on a 10 mm square coolbase and 50 mm square × 25mm high alpha heat sink and set under circular beam optic with integrated legs for parallel even light. The light pattern was programmed with BASIC Stamp 2.0 microcontroller and buckpuck DC driver (LUXEON, 700 mA, externally dimmable).

Local light stimulation was delivered by a 488 nm argon-ion laser using a Zeiss LSM 880 microscope. At 24 h AI, larvae were anesthetized and C4da neurons were imaged with a confocal microscope. For lesioned axons that bifurcated and formed two branches, we focused the laser beam (at 15% laser power) on the ventral branch for 5 min. The larva was then returned to grape juice agar plates and imaged again at 48 h AI to assess the increased length of each branch.

### Live cell imaging

For the light-induced membrane recruitment assay, BHK-21 cells were co-transfected with optoRaf or optoAKT. Fluorescence imaging of the transfected cells was carried out using a confocal microscope (Zeiss LSM 700). GFP fluorescence was excited by a 488-nm laser beam; mCherry fluorescence was excited by a 555-nm laser beam. Excitation beams were focused via a 40× oil objective (Plan-Neofluar NA 1.30). Ten pulsed 488-nm and 555-nm excitation were applied for each membrane recruitment experiment. CRY2-CIBN binding induced by 488-nm light was monitored by membrane recruitment of CRY2-mCherry-Raf1 (for optoRaf) or CRY2-mCherry-AKT (for optoAKT) to the CIBN-CIBN-GFP-CaaX-anchored plasma membrane. The powers after the objective for 488-nm and 555-nm laser beam are approximately 40 µW and 75 µW, respectively. Alternatively, an epi-illumination fluorescence microscope (Leica DMI8) equipped with a 100× objective (HCX PL FLUOTAR 100×/1.30 oil) and a light-emitting diode illuminator (SOLA SE II 365) was used for the CRY2-mCherry-Raf1 membrane translocation assay. Neurite outgrowth of PC12 cells was imaged using an epi-illumination fluorescence microscope (Leica DMI8) equipped with 10× (PLAN 10×/0.25) and 40× (HCXPL FL L 40×/0.6) objectives. Fluorescence from GFP was detected using the GFP filter cube (Leica, excitation filter 472/30, dichroic mirror 495, and emission filter 520/35); fluorescence from mCherry was detected using the Texas Red filter cube (Leica, excitation filter 560/40, dichroic mirror 595, and emission filter 645/75).

### Western Blot

Cells were washed once with 1 mL cold DPBS and lysed with 100 µL cold lysis buffer (RIPA + protease/phosphatase cocktail). Lysates were centrifuged at 17,000 RCF, 4 °C for 10 minutes to pellet cell debris. Purified lysates were normalized using Bradford reagent. Normalized samples were mixed with LDS buffer and loaded onto 10% or 12% polyacrylamide gels. SDS-PAGE was performed at room temperature with a cold water bath. Samples were transferred to PVDF membranes at 30 V 4 °C overnight or 80 V for 90 minutes. Membranes were blocked in 5% BSA/TBST for 1 hour at room temperature and probed with the primary and secondary antibodies according to company guidelines. Membranes were incubated with ECL substrate and imaged using a Bio-Rad ChemiDoc XRS chemiluminescence detector. Signal intensity analysis was performed by ImageJ.

### Statistical Analysis

No statistical methods were used to pre-determine sample sizes but our sample sizes are similar to those reported in previous publications (Song et al. 2012; Song et al. 2015), and the statistical analyses were done afterward without interim data analysis. Data distribution was assumed to be normal but this was not formally tested. All data were collected and processed randomly. Each experiment was successfully reproduced at least three times and was performed on different days. The values of ““*N*’’ (sample size) are provided in the figure legends. Data are expressed as mean ± SEM in bar graphs, if not mentioned otherwise. No data points were excluded. Two-tailed unpaired Student’s t-test was performed for comparison between two groups of samples. One-way ANOVA followed by multiple comparison test was performed for comparisons among three or more groups of samples. Two-way ANOVA followed by multiple comparison test was performed for comparisons between two or more curves. Fisher’s exact test was used to compare the percentage. Statistical significance was assigned, \**P* < 0.05, \*\**P* < 0.01, \*\*\**P* < 0.001.

## Acknowledgments

The plasmid of FOXO3-EGFP was a gift from Prof. Anne Brunet at Stanford University. This work was supported by NIH grants 1R01NS107392 (Y.S.) and 1R01GM132438 (K.Z.).

## Author Contributions

Y.S., K.Z. and Q.W. conceived the experimental design. Q.W., H.F., F.L., S.S.S, and V.V.K. conducted the experiments. Q. W. and H. F. analyzed the data. Q.W., H.F., Y.S., and K.Z. prepared the manuscript and figures.

## Supplemental Data

**Supplementary Figure 1.**
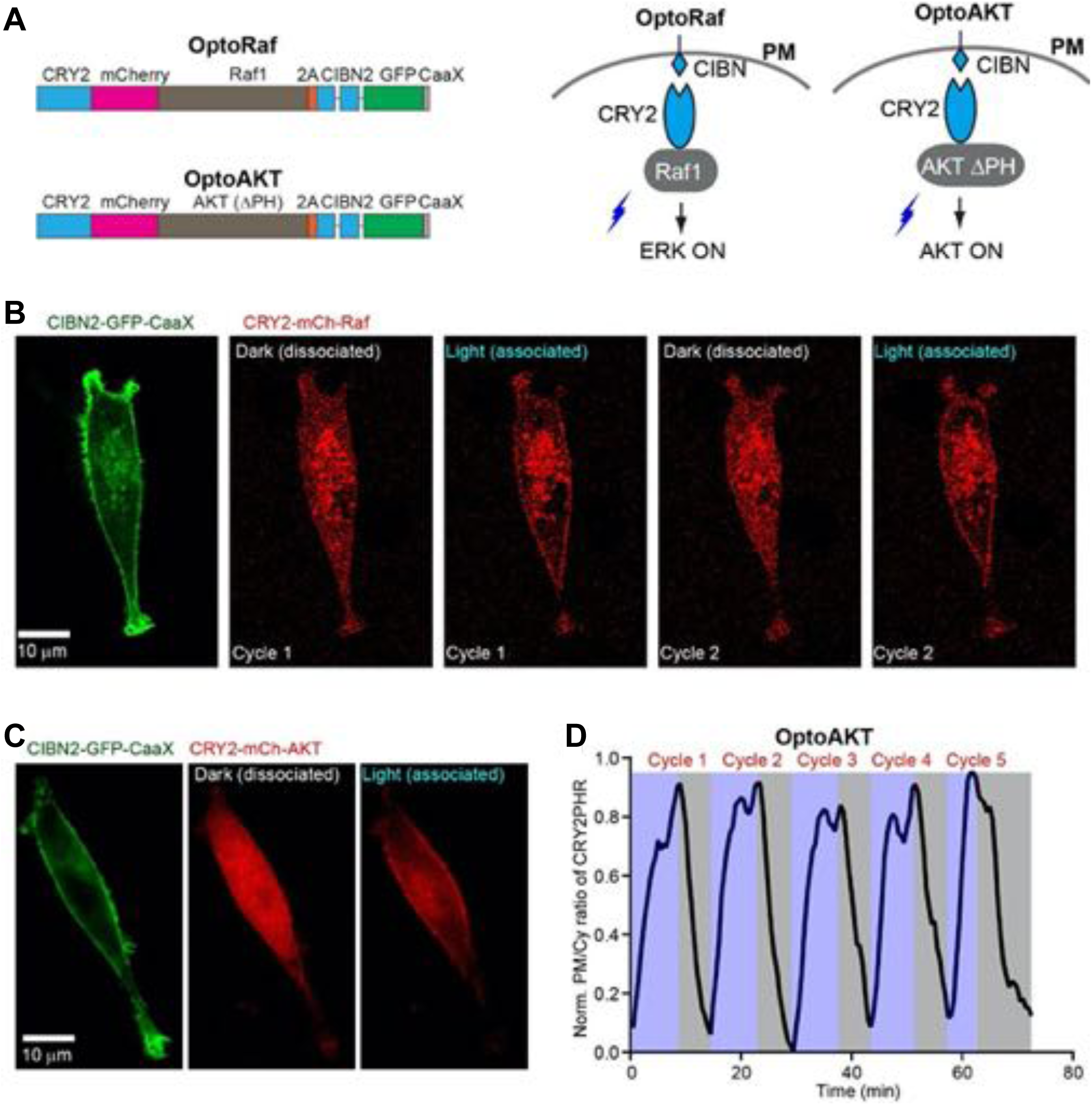
Design and live cell imaging for optoRaf and optoAKT in mammalian cell cultures. (*A*) Blue light illumination facilitates the association of CIBN and CRY2, and the CIBN/CRY2 complex spontaneously dissociates in the dark. In both optoRaf and optoAKT, CIBN-GFP-CaaX anchors to the plasma membrane and the cytosolic signaling protein was fused to CRY2. Under blue light stimulation, optoRaf and optoAKT recruit the signaling protein, Raf1 (optoRaf) and AKT LPH (optoAKT) to the plasma membrane to activate the ERK and AKT signaling pathway, respectively. (*B, C*) Live-cell imaging of reversible membrane recruitment of CRY2-mCh-Raf (B) CRY2-mCh-AKT (C). After each cycle of light stimulation, cells were kept in the dark for about 30 min. (*D*) Multiple cycles of membrane recruitment can be achieved from the same cell.

**Supplementary Figure 2.**
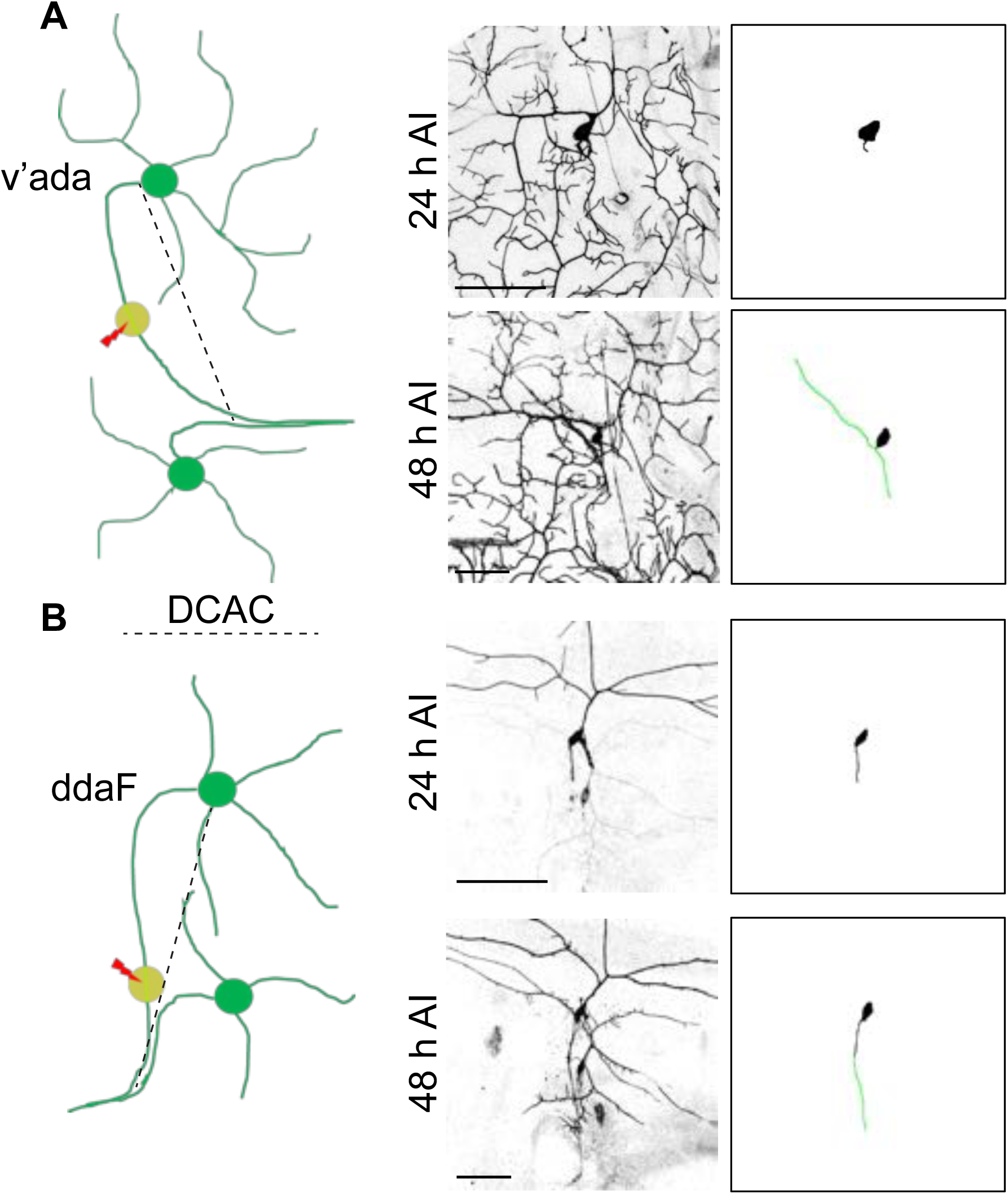
Quantification of axon regeneration in the PNS. (*A*) A schematic diagram depicts the C4da neuron injury model. At 48 h AI, two branches of the regenerating axon are extended towards two opposite directions. To calculate the regeneration index, the increased length of the longer branch was measured and normalized by DCAC (the distance between the cell body and the axon converging point). Scale bar = 50 µm. (*B*) A schematic drawing depicts the C3da neuron injury model. The green line depicts the regenerated axon. Scale bar = 50 µm. Related to **Figure. 3**.

**Supplementary Figure 3.**
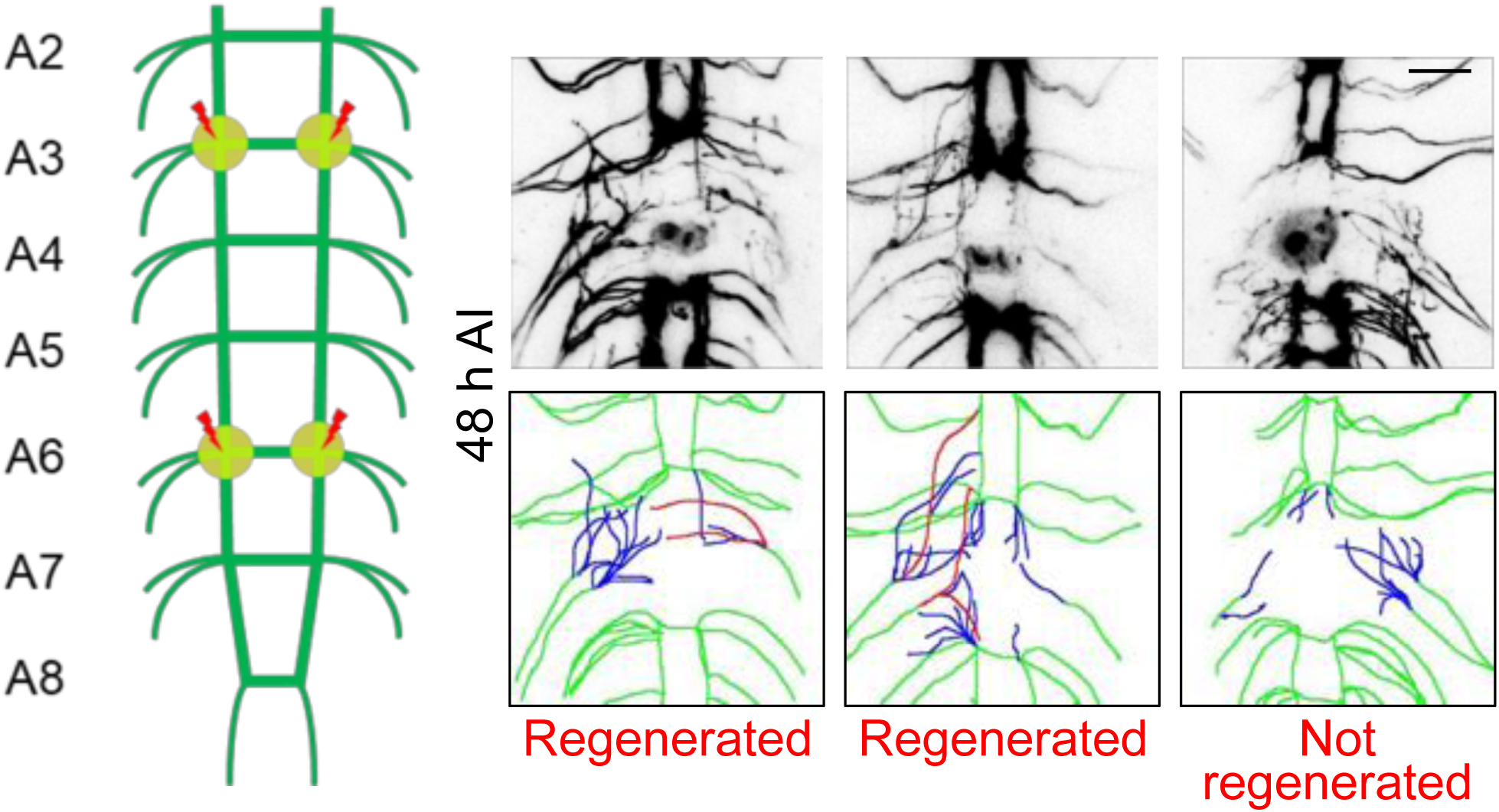
Quantification of axon regeneration in the CNS. A schematic diagram of the VNC injury method. The abdominal A3 and A6 bundles were injured by laser and the regeneration of these two commissure segments were assessed independently at 48 h AI. The regeneration index is defined as the total length of all regenerated axons normalized to the length between A4 and A5 bundles. However, a commissure segment is defined as regenerated only when at least one axon extends beyond the midline of the commissure region or connects with other intact bundles. Those axons are illustrated in the schematic diagrams in red, while other regrowing axons are in blue. Scale bar = 20 µm. Related to **Figure. 5**.

Supplementary movies of reversible optogenetic stimulation of Raf and AKT membrane recruitment, nuclear translocation of ERK, nuclear retreatment of FOXO3-GFP with optoRaf resolved by live-cell imaging:

**Movie S1:** Reversible optogenetic stimulation of Raf membrane recruitment with optoRaf resolved by live-cell imaging in BHK21 cells.

**Movie S2:** Reversible optogenetic stimulation of AKT membrane recruitment with optoAKT resolved by live-cell imaging in BHK21 cells.

**Movie S3:** Optogenetic activation of optoRaf causes nuclear translocation of ERK-GFP in BHK21 cells.

**Movie S4:** Optogenetic activation of optoAKT causes retreatment of FOXO3-GFP from the nucleus into the cytoplasm in BHK21 cells.

## References

Aloe L, Rocco ML, Bianchi P, Manni L. 2012. Nerve growth factor: from the early discoveries to the potential clinical use. J Transl Med 10.

Awasaki T, Lai SL, Ito K, Lee T. 2008. Organization and postembryonic development of glial cells in the adult central brain of Drosophila. J Neurosci 28: 13742–13753.

Bibel M, Barde YA. 2000. Neurotrophins: key regulators of cell fate and cell shape in the vertebrate nervous system. Genes Dev 14: 2919–2937.

Cervellini I, Galino J, Zhu N, Allen S, Birchmeier C, Bennett DL. 2018. Sustained MAPK/ERK Activation in Adult Schwann Cells Impairs Nerve Repair. J Neurosci 38: 679–690.

Chao MV. 2003. Neurotrophins and their receptors: A convergence point for many signalling pathways. Nature Reviews Neuroscience 4: 299–309.

Chen S, Weitemier AZ, Zeng X, He L, Wang X, Tao Y, Huang AJY, Hashimotodani Y, Kano M, Iwasaki H et al. 2018. Near-infrared deep brain stimulation via upconversion nanoparticle-mediated optogenetics. Science 359: 679–684.

Cheng PL, Song AH, Wong YH, Wang S, Zhang X, Poo MM. 2011. Self-amplifying autocrine actions of BDNF in axon development. Proc Natl Acad Sci U S A 108: 18430–18435.

Cho HJ, Kim JK, Park HC, Kim JK, Kim DS, Ha SO, Hong HS. 1998. Changes in brain-derived neurotrophic factor immunoreactivity in rat dorsal root ganglia, spinal cord, and gracile nuclei following cut or crush injuries. Exp Neurol 154: 224–230.

Dagliyan O, Hahn KM. 2019. Controlling protein conformation with light. Curr Opin Struct Biol 57: 17–22.

Dine E, Gil AA, Uribe G, Brangwynne CP, Toettcher JE. 2018. Protein Phase Separation Provides Long-Term Memory of Transient Spatial Stimuli. Cell Syst 6: 655–663.

Emoto K, Parrish JZ, Jan LY, Jan YN. 2006. The tumour suppressor Hippo acts with the NDR kinases in dendritic tiling and maintenance. Nature 443: 210–213.

Fang H, Liu C, Yang M, Li H, Zhang F, Zhang W, Zhang J. 2017. Neurotrophic factor and Trk signaling mechanisms underlying the promotion of motor recovery after acute spinal cord injury in rats. Exp Ther Med 14: 652–656.

Fukuoka T, Kondo E, Dai Y, Hashimoto N, Noguchi K. 2001. Brain-derived neurotrophic factor increases in the uninjured dorsal root ganglion neurons in selective spinal nerve ligation model. J Neurosci 21: 4891–4900.

Gao FB, Brenman JE, Jan LY, Jan YN. 1999. Genes regulating dendritic outgrowth, branching, and routing in Drosophila. Genes & development 13: 2549–2561.

Goglia AG, Toettcher JE. 2019. A bright future: optogenetics to dissect the spatiotemporal control of cell behavior. Curr Opin Chem Biol 48: 106–113.

Grueber WB, Jan LY, Jan YN. 2002. Tiling of the Drosophila epidermis by multidendritic sensory neurons. Development 129: 2867–2878.

Guo X, Snider WD, Chen B. 2016. GSK3beta regulates AKT-induced central nervous system axon regeneration via an eIF2Bepsilon-dependent, mTORC1-independent pathway. Elife 5: e11903.

Han C, Jan LY, Jan YN. 2011. Enhancer-driven membrane markers for analysis of nonautonomous mechanisms reveal neuron-glia interactions in Drosophila. Proc Natl Acad Sci U S A 108: 9673–9678.

Hayashi M, Ueyama T, Nemoto K, Tamaki T, Senba E. 2000. Sequential mRNA expression for immediate early genes, cytokines, and neurotrophins in spinal cord injury. J Neurotrauma 17: 203–218.

He L, Zhang YW, Ma GL, Tan P, Li ZJ, Zang SB, Wu X, Jing J, Fang SH, Zhou LJ et al. 2015. Near-infrared photoactivatable control of Ca2+ signaling and optogenetic immunomodulation. Elife 4.

He Z, Jin Y. 2016. Intrinsic Control of Axon Regeneration. Neuron 90: 437–451.

Huang EJ, Reichardt LF. 2001. Neurotrophins: roles in neuronal development and function. Annu Rev Neurosci 24: 677–736.

Huang H, Liu H, Yan R, Hu M. 2017. PI3K/Akt and ERK/MAPK Signaling Promote Different Aspects of Neuron Survival and Axonal Regrowth Following Rat Facial Nerve Axotomy. Neurochem Res 42: 3515–3524.

Hyung S, Lee SR, Kim YJ, Bang S, Tahk D, Park JC, Suh JF, Jeon NL. 2019. Optogenetic neuronal stimulation promotes axon outgrowth and myelination of motor neurons in a three-dimensional motor neuron-Schwann cell coculture model on a microfluidic biochip. Biotechnol Bioeng 116: 2425–2438.

Jeong JW, McCall JG, Shin G, Zhang Y, Al-Hasani R, Kim M, Li S, Sim JY, Jang KI, Shi Y et al. 2015. Wireless Optofluidic Systems for Programmable In Vivo Pharmacology and Optogenetics. Cell 162: 662–674.

Johnson HE, Toettcher JE. 2018. Illuminating developmental biology with cellular optogenetics. Curr Opin Biotechnol 52: 42–48.

Joo W, Hippenmeyer S, Luo L. 2014. Neurodevelopment. Dendrite morphogenesis depends on relative levels of NT-3/TrkC signaling. Science 346: 626–629.

Kaneko T, Macara AM, Li R, Hu Y, Iwasaki K, Dunnings Z, Firestone E, Horvatic S, Guntur A, Shafer OT et al. 2017. Serotonergic Modulation Enables Pathway-Specific Plasticity in a Developing Sensory Circuit in Drosophila. Neuron 95: 623–638.

Kennedy MJ, Hughes RM, Peteya LA, Schwartz JW, Ehlers MD, Tucker CL. 2010. Rapid blue-light-mediated induction of protein interactions in living cells. Nat Methods 7: 973–975.

Khamo JS, Krishnamurthy VV, Chen Q, Diao J, Zhang K. 2019. Optogenetic Delineation of Receptor Tyrosine Kinase Subcircuits in PC12 Cell Differentiation. Cell Chem Biol 26: 400–410

Khamo JS, Krishnamurthy VV, Sharum SR, Mondal P, Zhang K. 2017. Applications of Optobiology in Intact Cells and Multicellular Organisms. J Mol Biol 429: 2999–3017.

Krishnamurthy VV, Khamo JS, Mei W, Turgeon AJ, Ashraf HM, Mondal P, Patel DB, Risner N, Cho EE, Yang J et al. 2016. Reversible optogenetic control of kinase activity during differentiation and embryonic development. Development 143: 4085–4094.

Kuo CT, Jan LY, Jan YN. 2005. Dendrite-specific remodeling of Drosophila sensory neurons requires matrix metalloproteases, ubiquitin-proteasome, and ecdysone signaling. Proceedings of the National Academy of Sciences of the United States of America 102: 15230–15235.

Kuo CT, Zhu S, Younger S, Jan LY, Jan YN. 2006. Identification of E2/E3 ubiquitinating enzymes and caspase activity regulating Drosophila sensory neuron dendrite pruning. Neuron 51: 283–290.

Kyung T, Lee S, Kim JE, Cho T, Park H, Jeong YM, Kim D, Shin A, Kim S, Baek J et al. 2015. Optogenetic control of endogenous Ca(2+) channels in vivo. Nat Biotechnol 33: 1092–1096.

Leopold AV, Chernov KG, Verkhusha VV. 2018. Optogenetically controlled protein kinases for regulation of cellular signaling. Chem Soc Rev.

Liu BP, Cafferty WB, Budel SO, Strittmatter SM. 2006. Extracellular regulators of axonal growth in the adult central nervous system. Philos Trans R Soc Lond B Biol Sci 361: 1593–1610.

Liu K, Tedeschi A, Park KK, He Z. 2011. Neuronal intrinsic mechanisms of axon regeneration. Annu Rev Neurosci 34: 131–152.

Lizcano JM, Alrubaie S, Kieloch A, Deak M, Leevers SJ, Alessi DR. 2003. Insulin-induced Drosophila S6 kinase activation requires phosphoinositide 3-kinase and protein kinase B. Biochem J 374: 297–306.

Lu Y, Belin S, He Z. 2014. Signaling regulations of neuronal regenerative ability. Curr Opin Neurobiol 27: 135–142.

Ma G, Liu J, Ke Y, Liu X, Li M, Wang F, Han G, Huang Y, Wang Y, Zhou Y. 2018. Optogenetic Control of Voltage-Gated Calcium Channels. Angew Chem Int Ed Engl 57: 7019–7022.

Mahar M, Cavalli V. 2018. Intrinsic mechanisms of neuronal axon regeneration. Nat Rev Neurosci 19: 323–337.

Markus A, Zhong J, Snider WD. 2002. Raf and akt mediate distinct aspects of sensory axon growth. Neuron 35: 65–76.

Marshall CJ. 1995. Specificity of receptor tyrosine kinase signaling: transient versus sustained extracellular signal-regulated kinase activation. Cell 80: 179–185.

Miao L, Yang L, Huang H, Liang F, Ling C, Hu Y. 2016. mTORC1 is necessary but mTORC2 and GSK3beta are inhibitory for AKT3-induced axon regeneration in the central nervous system. Elife 5: e14908.

Miron M, Lasko P, Sonenberg N. 2003. Signaling from Akt to FRAP/TOR targets both 4E-BP and S6K in Drosophila melanogaster. Mol Cell Biol 23: 9117–9126.

Mitre M, Mariga A, Chao MV. 2017. Neurotrophin signalling: novel insights into mechanisms and pathophysiology. Clin Sci 131: 13–23.

Motta-Mena LB, Reade A, Mallory MJ, Glantz S, Weiner OD, Lynch KW, Gardner KH. 2014. An optogenetic gene expression system with rapid activation and deactivation kinetics. Nat Chem Biol 10: 196–202.

O’Donovan KJ, Ma K, Guo H, Wang C, Sun F, Han SB, Kim H, Wong JK, Charron J, Zou H et al. 2014. B-RAF kinase drives developmental axon growth and promotes axon regeneration in the injured mature CNS. J Exp Med 211: 801–814.

Ong Q, Guo S, Duan L, Zhang K, Collier EA, Cui B. 2016. The Timing of Raf/ERK and AKT Activation in Protecting PC12 Cells against Oxidative Stress. PLoS One 11: e0153487.

Park KK, Liu K, Hu Y, Smith PD, Wang C, Cai B, Xu B, Connolly L, Kramvis I, Sahin M et al. 2008. Promoting axon regeneration in the adult CNS by modulation of the PTEN/mTOR pathway. Science 322: 963–966.

Park S, Koppes RA, Froriep UP, Jia X, Achyuta AK, McLaughlin BL, Anikeeva P. 2015a. Optogenetic control of nerve growth. Sci Rep 5: 9669.

Park SI, Brenner DS, Shin G, Morgan CD, Copits BA, Chung HU, Pullen MY, Noh KN, Davidson S, Oh SJ et al. 2015b. Soft, stretchable, fully implantable miniaturized optoelectronic systems for wireless optogenetics. Nat Biotechnol 33: 1280–1286.

Parrish JZ, Emoto K, Kim MD, Jan YN. 2007. Mechanisms that regulate establishment, maintenance, and remodeling of dendritic fields. Annu Rev Neurosci 30: 399–423.

Poyhonen S, Er S, Domanskyi A, Airavaara M. 2019. Effects of Neurotrophic Factors in Glial Cells in the Central Nervous System: Expression and Properties in Neurodegeneration and Injury. Front Physiol 10: 486.

Ramer MS, Priestley JV, McMahon SB. 2000. Functional regeneration of sensory axons into the adult spinal cord. Nature 403: 312–316.

Rose CR, Blum R, Pichler B, Lepier A, Kafitz KW, Konnerth A. 2003. Truncated TrkB-T1 mediates neurotrophin-evoked calcium signalling in glia cells. Nature 426: 74–78.

Schwab ME, Strittmatter SM. 2014. Nogo limits neural plasticity and recovery from injury. Curr Opin Neurobiol 27: 53–60.

Shin Y, Berry J, Pannucci N, Haataja MP, Toettcher JE, Brangwynne CP. 2017. Spatiotemporal Control of Intracellular Phase Transitions Using Light-Activated optoDroplets. Cell 168: 159–171 e114.

Silver J, Miller JH. 2004. Regeneration beyond the glial scar. Nat Rev Neurosci 5: 146–156.

Song Y, Ori-McKenney KM, Zheng Y, Han C, Jan LY, Jan YN. 2012. Regeneration of Drosophila sensory neuron axons and dendrites is regulated by the Akt pathway involving Pten and microRNA bantam. Genes Dev 26: 1612–1625.

Song Y, Sretavan D, Salegio EA, Berg J, Huang X, Cheng T, Xiong X, Meltzer S, Han C, Nguyen TT et al. 2015. Regulation of axon regeneration by the RNA repair and splicing pathway. Nat Neurosci 18: 817–825.

Stone MC, Albertson RM, Chen L, Rolls MM. 2014. Dendrite injury triggers DLK-independent regeneration. Cell reports 6: 247–253.

Sugimura K, Yamamoto M, Niwa R, Satoh D, Goto S, Taniguchi M, Hayashi S, Uemura T. 2003. Distinct developmental modes and lesion-induced reactions of dendrites of two classes of Drosophila sensory neurons. The Journal of neuroscience : the official journal of the Society for Neuroscience 23: 3752–3760.

Sun F, He Z. 2010. Neuronal intrinsic barriers for axon regeneration in the adult CNS. Curr Opin Neurobiol 20: 510–518.

Sun L, Shay J, McLoed M, Roodhouse K, Chung SH, Clark CM, Pirri JK, Alkema MJ, Gabel CV. 2014. Neuronal Regeneration in C. elegans Requires Subcellular Calcium Release by Ryanodine Receptor Channels and Can Be Enhanced by Optogenetic Stimulation. Journal of Neuroscience 34: 15947–15956.

Sung KJ, Yang WL, Wu CB. 2019. Uncoupling neurotrophic function from nociception of nerve growth factor: what can be learned from a rare human disease? Neural Regen Res 14: 570–573.

Wang W, Wildes CP, Pattarabanjird T, Sanchez MI, Glober GF, Matthews GA, Tye KM, Ting AY. 2017. A light- and calcium-gated transcription factor for imaging and manipulating activated neurons. Nat Biotechnol 35: 864–871.

Watson FL, Heerssen HM, Bhattacharyya A, Klesse L, Lin MZ, Segal RA. 2001. Neurotrophins use the Erk5 pathway to mediate a retrograde survival response. Nat Neurosci 4: 981–988.

Williams DW, Kondo S, Krzyzanowska A, Hiromi Y, Truman JW. 2006. Local caspase activity directs engulfment of dendrites during pruning. Nature neuroscience 9: 1234–1236.

Williams DW, Truman JW. 2005. Cellular mechanisms of dendrite pruning in Drosophila: insights from in vivo time-lapse of remodeling dendritic arborizing sensory neurons. Development 132: 3631–3642.

Wu X, Zhang Y, Takle K, Bilsel O, Li Z, Lee H, Zhang Z, Li D, Fan W, Duan C et al. 2016. Dye-Sensitized Core/Active Shell Upconversion Nanoparticles for Optogenetics and Bioimaging Applications. ACS Nano 10: 1060–1066.

Wu YI, Frey D, Lungu OI, Jaehrig A, Schlichting I, Kuhlman B, Hahn KM. 2009. A genetically encoded photoactivatable Rac controls the motility of living cells. Nature 461: 104–108.

Xiang Y, Yuan Q, Vogt N, Looger LL, Jan LY, Jan YN. 2010. Light-avoidance-mediating photoreceptors tile the Drosophila larval body wall. Nature 468: 921–926.

Xiao JH, Wong AW, Willingham MM, van den Buuse M, Kilpatrick TJ, Murray SS. 2010. Brain-Derived Neurotrophic Factor Promotes Central Nervous System Myelination via a Direct Effect upon Oligodendrocytes. Neurosignals 18: 186–202.

Xiao Y, Tian W, Lopez-Schier H. 2015. Optogenetic stimulation of neuronal repair. Curr Biol 25: R1068–1069.

Yiu G, He Z. 2006. Glial inhibition of CNS axon regeneration. Nat Rev Neurosci 7: 617–627.

Zhang K, Cui B. 2015. Optogenetic control of intracellular signaling pathways. Trends Biotechnol 33: 92–100.

Zhang K, Duan L, Ong Q, Lin Z, Varman PM, Sung K, Cui B. 2014. Light-mediated kinetic control reveals the temporal effect of the Raf/MEK/ERK pathway in PC12 cell neurite outgrowth. PLoS One 9: e92917.

Zhao EM, Suek N, Wilson MZ, Dine E, Pannucci NL, Gitai Z, Avalos JL, Toettcher JE. 2019. Light-based control of metabolic flux through assembly of synthetic organelles. Nat Chem Biol 15: 589–597.

Zhao EM, Zhang Y, Mehl J, Park H, Lalwani MA, Toettcher JE, Avalos JL. 2018. Optogenetic regulation of engineered cellular metabolism for microbial chemical production. Nature 555: 683–687.

Zhou XX, Fan LZ, Li P, Shen K, Lin MZ. 2017. Optical control of cell signaling by single-chain photoswitchable kinases. Science 355: 836–842.

